# Genotype-phenotype relationships in children with Copy Number Variants associated with high neuropsychiatric risk: Findings from the case-control IMAGINE-ID cohort in the United Kingdom

**DOI:** 10.1101/535708

**Authors:** Samuel J.R.A. Chawner, Michael Owen, Peter Holmans, Lucy Raymond, David Skuse, Jeremy Hall, Marianne van den Bree

## Abstract

**Background:** A variety of Copy Number Variants are associated with a high risk of neurodevelopmental and psychiatric disorders (ND-CNVs). We aimed to characterise the impact of ND-CNVs on childhood development and investigate whether different ND-CNVs lead to distinct and specific patterns of cognitive and behavioural outcomes.

**Methods:** 258 children with ND-CNVs (13 CNVs across 9 loci) were systematically assessed for psychiatric disorders as well as broader traits of neurodevelopmental, cognitive and psychopathological origin. A comparison was made with 106 control siblings, in order to test the hypothesis that phenotypes would differ by genotype, both quantitatively, in terms of severity, and qualitatively in the pattern of associated impairments.

**Outcomes:** 79.8% of ND-CNVs carriers met criteria for one or more psychiatric disorders (OR=13.8 compared to controls): the risk of ADHD (OR=6.9), ODD (OR=3.6), anxiety disorders (OR=2.9), and ASD traits (OR=44.1) was particularly high. ND-CNVs carriers were impaired across all neurodevelopmental, cognitive, and psychopathological traits relative to controls. Only moderate quantitative and qualitative differences in phenotypic profile were found between genotypes. In general, the range of phenotypes was broadly similar for all ND-CNV genotypes. Traits did show some evidence of genotypic specificity, however the specific genotype accounted for a low proportion of variance in outcome (5-20% depending on trait).

**Interpretation:** The 13 ND-CNVs studied have a similar range of adverse effects on childhood neurodevelopment, despite subtle quantitative and qualitative differences. Our findings suggest that genomic risk for neuropsychiatric disorder has pleiotropic effects on multiple processes and neural circuits, and provides important implications for research into genotype-phenotype relationships within psychiatry.

**Funding:** The Medical Research Council and the Waterloo Foundation

**Research in context:** *Evidence before this study:* Several Copy Number Variants (CNVs) have been associated with high risk of development of child and adult neuropsychiatric disorders. Increasingly young children with developmental delay referred for genetic testing are being diagnosed with neurodevelopmental and psychiatric risk CNVs (referred to as ND-CNVs hereafter). It remains unclear whether different genotypes are associated with specific cognitive and behavioural phenotypes or whether these outcomes are non-specific. We searched PubMed for English language studies published from database inception until January 10th, 2019 that investigated the relationship between CNVs and cognitive and behavioural outcomes. Search terms included “CNV”, “genomics”, “1q21.1”, “2p16.3”, “NRXN1”, “9q34”, “Kleefstra Syndrome”, “15q11.2”, “15q13.3”, “16p11.2”, “22q11.2”, “psychiatry”, and “cognition”. Preliminary studies have indicated that deletions and duplications at the same loci may differ in cognitive and behavioural phenotypes. However, to date, there have been limited studies that contrasted the phenotypes of ND-CNVs across several loci on a range of cognitive and behavioural domains.

*Added value of this study:* We found that young people carrying a ND-CNV were at considerably increased risk for neuropsychiatric disorder and impairments across a range of neurodevelopmental, psychopathological, cognitive, social, sleep and motor traits. Within ND-CNV carriers, comparisons between genotypes indicated moderate quantitative and qualitative differences in overall phenotypic profile, with evidence that severity of impairment was similar across all genotypes for some traits (e.g. mood problems, sleep impairments, peer problems, and sustained attention) whereas for other traits there was evidence of genotype specific effects on severity (e.g., IQ, spatial planning, processing speed, subclinical psychotic experiences, ASD traits, motor coordination total psychiatric symptomatology, particularly anxiety, ADHD, and conduct related traits). However the proportion of variance explained by genotype was low, 5-20% depending on trait, indicating that overall ND-CNVs lead to similar neurodevelopmental outcomes. It is important that genotype-phenotype relationships are viewed through a developmental lens as some phenotypic outcomes were found to be associated with age.

*Implications of all the available evidence:* Our work highlights that children who carry a ND-CNV represent a patient group that warrants clinical and educational attention for a broad range of cognitive and behavioural impairments and that commonalities in clinically relevant neurodevelopmental impairments exist across ND-CNVs. This group of young people could benefit from the development of a general care pathway, to which genotype-specific recommendations can be added where needed. Our work indicates that the relationship between genotype and neurodevelopmental phenotype is complex and that future research will need to take a global systems approach and not be narrowly focused on single phenotypes.

## Introduction

The advent of microarray technology has heralded a new era for understanding the clinical genetics of neuropsychiatric disorders. A striking finding has been the implication of copy number variants (CNVs) in these disorders ^1^, including intellectual disability (ID), autism spectrum disorder (ASD) and schizophrenia ^2-4^. CNVs are submicroscopic deletions or duplications within the genome that are greater than 1000 base pairs^5^ and several loci have been identified whereby CNVs recur with sufficient frequency in the population to be associated with neurodevelopmental and psychiatric outcomes (hereafter referred to as ND-CNVs). Recurrent ND-CNVs are individually rare, but collectively pathogenic ND-CNVs have been implicated in ∼15% of patients with neurodevelopmental disability ^6^. Although these ND-CNVs are strongly associated with disorder, they have incomplete penetrance and exhibit a high degree of pleiotropy, conferring risk for a broad range of psychiatric disorders, cognitive deficits and medical/physical comorbidities across the lifespan ^7-10^.

Current understanding of genotype-phenotype relationships is hampered by a lack of studies that have conducted cross-CNV comparisons^11^. Therefore it is unclear to what extent phenotypic findings for different genotypes can be compared across studies and what the impact is of variation in sample sizes and methodological issues of ascertainment and phenotyping. Increasing use of array screening in the assessment of children with neurodevelopmental delay is leading to a rise in the diagnosis of ND-CNVs by medical genetics clinics, yet information on long-term neuropsychiatric prognosis is lacking. There is a need to understand whether different genotypes are associated with specific neuropsychiatric, cognitive and other phenotypes. We posit four different models of potential genotype-phenotype relationships (Figure 1). 1) The null model proposes that phenotypic profile does not differ between genotypes (Model 1, Figure 1). 2) Phenotypic differences are qualitative in nature, whereby each ND-CNV is associated with a distinct phenotype due to the specific genes involved (Model 2, Figure 1). 3) Phenotypic differences are quantitative in nature whereby all ND-CNVs impact on the same range of outcomes but differ from each other in magnitude of impairment (Model 3, Figure 1). 4) A combination of the Models 2 and 3 best explains differences in phenotypic outcome across ND-CNVs (Model 4, Figure1). There is support for the qualitative differences model in the autism field where it is hypothesised that the disorder is dissociable by the genetic underpinnings^12,13^, with some researchers using the term “autisms”^14^. The quantitative differences model is supported by findings that genes across ND-CNVs impact shared pathways leading to outcomes such as cognitive impairment^15^ and increased schizophrenia risk^16^, indicating that common mechanisms act across loci. It is important to highlight that variability in phenotypic outcomes will also be shaped by incomplete penetrance^8^, life course developmental stage^17^, genetic context including polygenic risk^18^ and additional mutations ^19,20^ as well as environmental exposures^21^.

**Figure 1:**
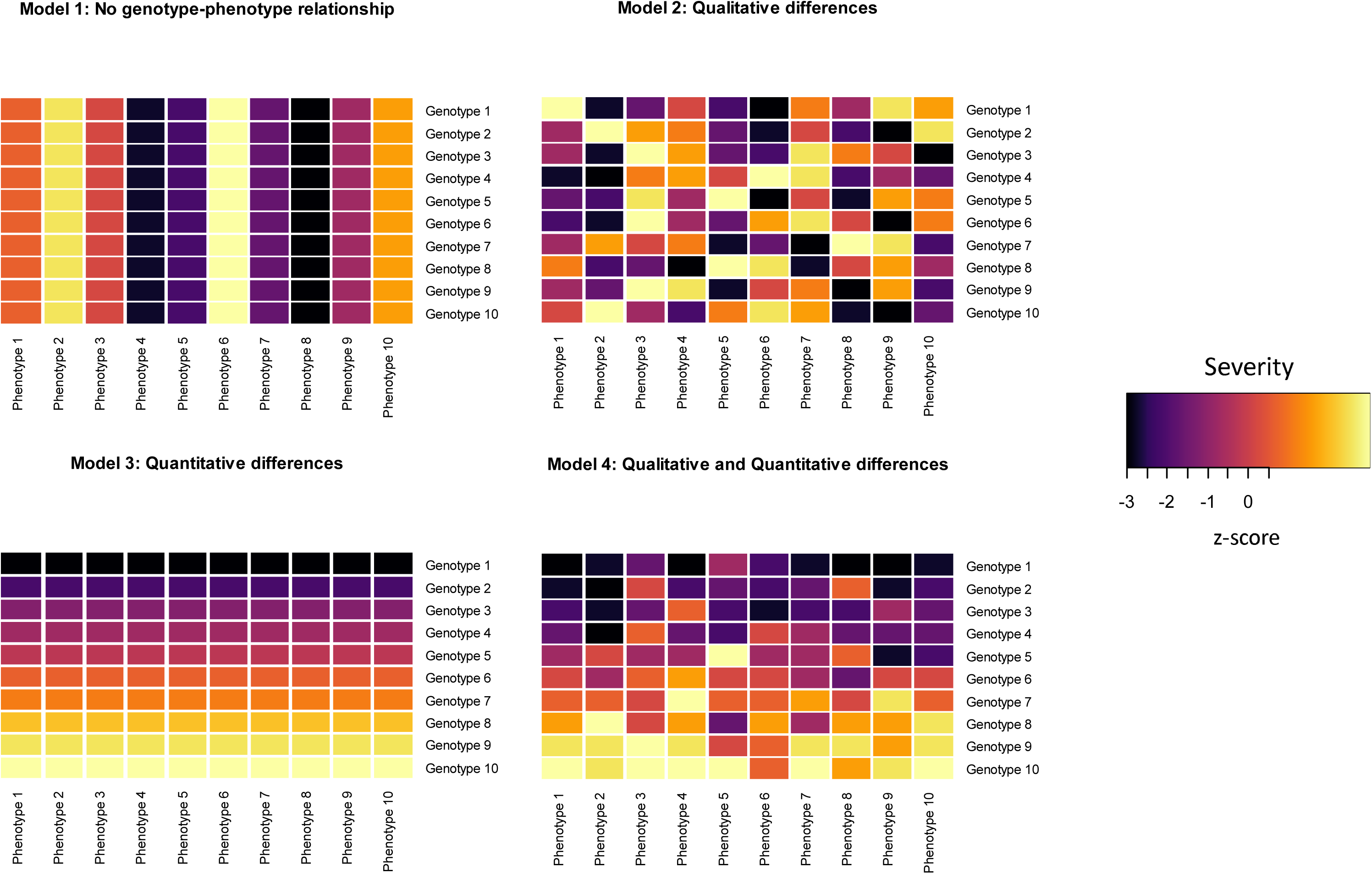
Visual representation of models of genotype-phenotype relationships. Each cell represents z-score performance for a neuropsychiatric domain. Z scores can be found in Supplementary Table 1. In model 1 there are no phenotypic differences between genotypes. In model 2 there are qualitative differences, in that the neuropsychiatric profile differs by genotype, but they are no quantitative differences. Each genotype has the same overall severity of impairment, but the distribution across phenotypic traits is different, e.g. for genotype 1, phenotype 2 and 6 are most severely affected, whilst for genotype 3 it is phenotypes 2 and 10. In model 3 there are quantitative differences as each genotype differs in average level of impairment, however, there are no qualitative differences in phenotypic profile as within each genotype, severity does not differ by phenotype. In model 4 there are both quantitative and qualitative differences in neuropsychiatric profile.

Here we present findings from a cohort of children with ND-CNVs from the IMAGINE-ID (Intellectual Disability & Mental Health: Assessing the Genomic Impact on Neurodevelopment) consortium. Individuals were recruited on the basis of genotype via the United Kingdom’s (UK) National Health Service (NHS) medical genetic clinic network. Broad online phenotyping was conducted on over 2000 individuals (results will be reported elsewhere), and deep phenotyping was conducted within a subgroup of the cohort with assessments covering a range of neuropsychiatric, cognitive and other traits using a multi-informant approach. The IMAGINE-ID study is creating a lasting resource for research into intellectual disability, genotype and mental health. Researchers who would like to use IMAGINE-ID data can find further information on our website “http://imagine-id.org/healthcare-professionals/. Here we report on findings from the deep phenotyping component of IMAGINE-ID. First, we characterised the impact of recurrent ND-CNVs on child development by contrasting the performance of CNV carriers with sibling controls. Next, we evaluated the phenotypic differences between genotypes and determined whether these were qualitative or quantitative in nature. Finally, we established the extent to which neuropsychiatric, cognitive and other outcomes were affected by gender and age.

## Methods

The Imagine-ID study recruits individuals with genomic variants that have been associated with neurodevelopmental problems. Participants are recruited via the UK NHS medical genetics service, whereby microarray results can be accessed and patients can be retrospectively and prospectively invited to take part in research studies. NHS patients were also recruited via support groups, including Unique, Max Appeal and other groups on social media. The IMAGINE-ID study comprises two components. First, parents with a child aged 4 years and older with a CNV or Single nucleotide polymorphisms (SNV) were invited to complete relatively short online assessments (>2000 completed to date). Second, from this pool, families with a child with one of a set of specific, recurrent ND-CNVs were approached for a deep phenotyping home based assessment and it is this sample that the current study is based on. The specific loci were: 1q21.1 (proximal and distal), 2p16.3, 9q34, 15q11.2, 15q13.3, 16p11.2 (proximal and distal) and 22q11.2 (Table 1 for further details). These recurrent ND-CNVs were selected because they are robustly associated with ID and neuropsychiatric phenotypes^22-24^, including schizophrenia and ASD, and are frequently diagnosed in medical genetic clinics. Families were approached if the child with the ND-CNV was aged 6-19 years (the age range in which our assessment battery operates) and if presence of the ND-CNV was confirmed from accessing medical records. 274 children with one of these ND-CNVs took part in these detailed assessments. 16 were found to have more than one of these ND-CNVs and were excluded from the analysis. This left a sample of 258 children with a ND-CNV (9.7 years (SD=3.1), 65.9% male). Of the 258 ND-CNV carriers 22.8% (n=59) had a de novo variant, 44.2% (n=114) an inherited variant and for 32.9% (n=85) the status was unknown. A sibling without the ND-CNV (sibling control) and closest in age to the index child was also invited to take part. We recruited 106 sibling controls (10.9 years (SD=3.0), 51.9% male)). Availability of micro-array results and medical records allowed us to exclude presence of the ND-CNVs under study for n=77. For the remaining, n=16 came from families with an inherited ND-CNV. Informed consent was gained from primary carers and participants. Protocols were approved by the NHS London Queen Square research ethics committee. ND-CNV genotype was confirmed via NHS medical genetics clinic records and by the Cardiff University Division of Psychological Medicine and Clinical Neurosciences (CU DPMCN) laboratory.

**Table 1:** ND-CNV breakpoints and frequencies # Not listed on Mendelian Inheritance in Man (MIM) website but associated with neurodevelopmental phenotypes.

| ND-CNV | Locus | Critical/Unique Sequence Region (hg19) | Gene/Locus<br>MIM<br>number | N |
| --- | --- | --- | --- | --- |
| 1q21.1 proximal duplication | 1q21.1 | chr1:145,394,955-145,807,817 | # | 14 |
| 1q21.1 distal deletion | 1q21.1 | chr1:146,527,987-147,394,444 | 612474 | 21 |
| 1q21.1 distal duplication | 1q21.1 | chr1:146,527,987-147,394,444 | 612475 | 21 |
| 2p16.3 deletion (*NRXN1*) | 2p16.3 | chr2:50145643-51259674 | 600565 | 14 |
| 9q34 del (Kleefstra, *EHMT1*) | 9q34.4 | chr9:140,513,444-140,730,578 | 607001 | 10 |
| 15q11.2 deletion BP1-BP2 | 15q11.2 | chr15:22,805,313-23,094,530 | 615656 | 35 |
| 15q13.3 deletion (*CHRNA7*) | 15q13.3 | chr15:32,017,070-32,453,068 | 612001 | 20 |
| 15q13.3 duplication (*CHRNA7*) | 15q13.3 | chr15:32,017,070-32,453,068 | # | 12 |
| 16p11.2 distal deletion (220kb) | 16p11.2 | chr16:28,823,196-29,046,783 | 613444 | 12 |
| 16p11.2 proximal deletion (593kb) | 16p11.2 | chr16:29,650,840-30,200,773 | 611913 | 44 |
| 16p11.2 proximal duplication (593kb) | 16p11.2 | chr16:29,650,840-30,200,773 | 614671 | 19 |
| 22q11.2 deletion (VCFS/DiGeorge) | 22q11.2 | chr22:19,037,332-21,466,726 | 602054 | 17 |
| 22q11.2 duplication | 22q11.2 | chr22:19,037,332-21,466,726 | 608363 | 19 |

## Assessments

25 quantitative cognitive and behavioural traits and 5 composite scores were measured using a multi-informant approach. In addition categorical psychiatric diagnoses were derived. Assessments of the child were made by experienced research psychologists. Assessments took place within the participant’s home with the advantages this maximised accessibility to the study and reduced bias against participants who may struggle to travel to a research clinic, and furthermore the child could be assessed in a familiar setting where they are less likely to be anxious and more likely to engage with the assessments. Measures are briefly described, full details on assessments and a summary table can be found in the Supplementary Materials.

### Cognition

Cognition was assessed via direct child assessments. IQ was assessed using the Wechsler Abbreviated Scale of Intelligence (WASI)^25^ from which scores for non-verbal reasoning, perceptual organisation, verbal knowledge and verbal reasoning were derived as well as full scale IQ (FSIQ), performance IQ (PIQ) and verbal IQ (VIQ) composite scores. Set-shifting ability was assessed using the Wisconsin Card Sorting Test (WSCT)^26^. The CANTAB (Cambridge Neuropsychological Test Automated Battery)^27^ was used to assess spatial working memory, spatial planning, sustained attention and processing speed.

### Psychopathology and functioning

The Child and Adolescent Psychiatric Assessment (CAPA)^28^ carer report interview was used to derive categorical diagnoses and a total symptom count composite score, as well as the following symptom subscales: attention deficit hyperactivity disorder (ADHD), anxiety, mood, obsessive-compulsive disorder (OCD), oppositional defiant disorder (ODD), and problems with sleep. The child report CAPA was conducted to assess subclinical psychotic experiences. Interviews were taped and diagnoses confirmed in consensus meeting led by a child psychiatrist. General and social functioning was assessed by the psychologists conducting the home visit using the Children’s Global Assessment Scale (CGAS)^29^ and the Social and Occupational Functioning Assessment Scale (SOFAS)^30^. Autism Spectrum Disorder (ASD) traits were assessed via caregiver report using the Social Communication Questionnaire (SCQ)^31^. Motor coordination impairment was assessed via caregiver report using the Developmental Coordination Disorder Questionnaire (DCDQ)^32^. The Strengths and Difficulties Questionnaire (SDQ)^33^ was completed by the caregiver and the teacher from which conduct, emotional, hyperactivity, peer (quality of peer relationships) and prosocial subscale scores were derived as well as SDQ total composite score.

## Analysis

### Aim 1: Cognitive and behavioural phenotype of ND-CNV carriers in relation to controls

#### Categorical outcome measures

The prevalence of psychiatric disorder was compared between ND-CNV carriers and controls. Analysis was conducted using generalized linear mixed-effects models, with carrier status, age and gender as fixed effects and family as a random effect to take into account that control siblings are related to ND-CNV carriers.

#### Continuous outcome measures

All cognitive and behavioural trait scores and composite scores (FSIQ, PIQ, VIQ, total symptom count and SDQ total score) were transformed using Tukey’s Ladder of Powers. This transformation makes the data fit the normal distribution as closely as possible. All tests scores were then standardised into z-scores using the mean and SDs of the control group as reference, i.e. the difference in the individual’s score and the mean score for the entire control group was divided by the SD for the control group. Z-scores were constructed so that a negative score denoted a poorer outcome. Linear mixed-effects models were conducted with test score as the outcome and carrier status, age and gender as fixed effects and family as a random effect. To estimate the standardised difference between ND-CNV carriers and controls Cohen’s d was calculated. To assess the potential effects of intelligence on group differences, analyses were repeated with FSIQ as a covariate. To correct for multiple testing in Aim 1 a Benjamini-Hochberg false discovery rate (B-H FDR) of 0.05 for correction of p-values was applied.

### Aim 2: Investigation of qualitative and quantitative differences between ND-CNVs

To investigate which genotype-phenotype relationship model (Figure 1) best explained our data, we used the z-scores generated within Aim 1 for each child with a ND-CNV to calculate the mean z-score across individuals with the same ND-CNV genotype for each trait. Hierarchical clustering was performed using Ward’s method and Euclidian distance to investigate which ND-CNVs, and which cognitive and behavioural phenotypes clustered together. The frequency of each ND-CNV is presented in Table 1.

Analysis of qualitative and quantitative differences in overall phenotypic profile was based on ranking the mean of each phenotypic trait score for each ND-CNV. In the analysis of qualitative effects a set of phenotype rankings was created for each ND-CNV (Model 2, Figure 1, within each genotype row, phenotype was ranked by phenotypic severity). Rank discordance between ND-CNVs would suggest that the phenotype profile for each ND-CNV differs, therefore indicating the presence of qualitative differences. In the analysis of quantitative effects a set of ND-CNV rankings was created for each phenotype (Model 3, Figure 1, within each phenotype column, each ND-CNV was ranked by phenotypic severity). Rank concordance between phenotypes would suggest that ND-CNVs differ in severity across phenotypes, therefore indicating quantitative differences. Note these models aren’t opposing ends of a spectrum, both quantitative and qualitative effects can be present (Model 4, Figure 1). To test for similarities and differences for both qualitative and quantitative effects across ND-CNVs, rank concordance was assessed using Kendall’s test and rank discordance using the Friedman test. This rank concordance based approach has been previously used to investigate genotype-phenotype relationships^34^. To avoid collinearity, composite scores were not included in the concordance analysis. Furthermore to test for quantitative effects between ND-CNVs at the level of individual traits, ANCOVAs were conducted with the test score as the outcome, genotype as the predictor, and gender and age as covariates.

### Aim 3: Effect of age and gender on cognitive and behavioural outcomes

To investigate the influence of gender and age on the outcomes within ND-CNV carriers, we estimated eta-squared and standardised beta value from the ANCOVAs conducted for the quantitative analysis. Eta-squared values reflect the proportion of variance in the quantitative trait explained by the predictor and the standardised beta values reflect the magnitude and direction of effect of the predictor on phenotypic outcome. To correct for analyses across aims 2 and 3 a B-H FDR 0.05 correction of p-values was applied.

## Results

*Aim 1:* Prevalence of psychiatric disorder was significantly elevated in ND-CNV carriers (79.8%) compared to controls (21.3%), OR = 13.8, 95% CI = 7.2-26.3, p=7.79×10^−7^). ND-CNV carriers had significantly elevated rates of ADHD (47.2% vs 11.0%, OR = 6.9, 95% CI = 3.2-15.1, 2.09×10^−6^), ODD (20.6% vs 6.7%, OR = 3.6, 95% CI = 1.4-9.4, p=1.20×10^−2^), anxiety disorder (21.9% vs 9.3%, OR = 2.9, 95% CI = 1.2-6.7, p=1.46×10^−2^), ASD (66.1% vs 4.7%, OR = 44.1, 95% CI = 15.3-127.5, 2.50×10^−9^) and Tic Disorder (16.3% vs 0.0%, p=2.10×10^−5^, OR could not be estimated as no controls affected) compared to controls (see Table 2). These results remained significant when FSIQ was controlled for, and all survived B-H FDR correction. Mood disorder, OCD and subclinical psychotic experiences were present in ND-CNV carriers but prevalence was not significantly elevated relative to controls. None of the ND-CNV carriers or controls met criteria for psychotic disorder.

**Table 2:** Prevalence of psychiatric disorder and childhood outcomes in ND-CNV carriers and controls CI, 95% confidence interval; OR, Odds ratio; ADHD, Attention Deficit Hyperactivity Disorder; ASD, Autism Spectrum Disorder; OCD, Obsessive Compulsive Disorder; ODD, Oppositional Defiant Disorder. Generalized linear mixed-effects model was conducted with diagnosis as the outcome and carrier status, age and gender as fixed effects and family as a random effect. #due to 0 values for controls OR could not be estimated, p-values were estimated using fishers exact test but should be treated cautiously *Survives Benjamini-Hochberg false discovery rate 0.05 correction

| Psychiatric diagnosis | ND-CNV carriers |  | Controls |  | ND-CNV carriers vs controls |  |  |
| --- | --- | --- | --- | --- | --- | --- | --- |
|  | N(%) | % CI | N(%) | % CI | p-value | OR | CI |
| Any Psychiatric Disorder | 186(79.8%) | 74.7,85.0 | 16(21.3%) | 12.1,30.6 | $7.79 \times 10^{-7} *$ | 13.8 | 7.2,26.3 |
| ADHD | 110(47.2%) | 40.8,53.6 | 8(11.0%) | 3.7,17.7 | $2.09 \times 10^{-6} *$ | 6.9 | 3.2,15.1 |
| Any Anxiety Disorder | 51(21.9%) | 16.6,27.2 | 7(9.3%) | 2.7,27.5 | $1.46 \times 10^{-2} *$ | 2.9 | 1.2,6.7 |
| Any Mood Disorder | 4(1.7%) | 0.0,3.4 | 1(1.3%) | -1.3,4.0 | $7.27 \times 10^{-1}$ | 1.5 | 0.2,14.0 |
| Any Psychotic Disorder | 0(0.0%) | - | 0(0.0%) | - | - | - | - |
| Subthreshold psychosis | 16(6.6%) | 4.1,10.4 | 1(1.2%) | 0.2,6.5 | $6.84 \times 10^{-2}$ | 6.7 | 0.9,52.2 |
| ASD | 150(66.1%) | 59.9,72.2 | 4(4.7%) | 0.0,9.1 | $2.50 \times 10^{-9} *$ | 44.1 | 15.3,127.5 |
| OCD | 8(3.4%) | 1.8,6.6 | 0(0.0%) | - | $2.06 \times 10^{-1} \#$ | - | - |
| ODD | 48(20.6%) | 15.4,25.8 | 5(6.7%) | 1.0,12.3 | $1.20 \times 10^{-2} *$ | 3.6 | 1.4,9.4 |
| Tic Disorders | 38(16.3%) | 12.1,21.6 | 0(0.0%) | - | $2.19 \times 10^{-5} * \#$ | - | - |

Linear mixed-effects model analysis found that ND-CNV carriers showed significant impairment on all our continuous measures of cognitive and behavioural traits and composite scores (Table 3, Figure 2) compared to controls. These results remained significant when FSIQ was controlled for, and all survived B-H FDR correction. Cohen’s d varied from 0.27 (subclinical psychotic experiences) to 1.76 (hyperactivity subscale, caregiver reported). Large effect size differences between ND-CNV carriers and controls were found for FSIQ, including PIQ, VIQ and all comprising subtests, sustained attention, total psychiatric symptom count, ADHD and ASD traits, motor coordination, general and social functioning, total SDQ score (carer and teacher report) and hyperactivity (carer and teacher report), peer (carer report), prosocial (carer report) SDQ subscales. The majority of contrasts remained significant when we compared all deletion carriers to the controls and all the duplications carriers to the controls (see Supplementary Table 1, which also details contrasts of each ND-CNV from controls). Findings remained similar when the 29 control sibling for whom we did not have full genetic confirmation of absence of the ND-CNVs under study were excluded (see Supplementary Table 2), therefore all control siblings were included in all subsequent analyses in this paper. As well as caregiver report, teachers reported that ND-CNV carriers scored significantly worse on SDQ total score and subscale scores (Table 3 and Figure 2). Teacher reported SDQ scores were moderately correlated with carer report scores; total SDQ score r = 0.470, p=1.35×10^−11^; SDQ subscale scores, r = 0.316 to 0.548, p = 5.69×10^−16^ to 1.13×10^−5^.

**Table 3:** Quantitative cognitive and behavioural traits in controls and ND-CNV carriers FSIQ, Full Scale Intelligence Quotient; PIQ, Performance Intelligence Quotient; VIQ, Verbal Intelligence Quotient; CAPA, Child and Adolescent Psychiatric Assessment; ADHD, Attention Deficit Hyperactivity Disorder; OCD, Obsessive Compulsive Disorder; ODD, Oppositional Defiant Disorder; ASD, Autism Spectrum Disorder; SDQ, Strengths and Difficulties Questionnaire. Test scores were transformed so that their distribution approximated the normal distribution as closely as possible. Transformed scores were standardised into z scores using the means and SDs of the control group as reference and adjusted for age and gender, and were constructed so that a negative score denoted a poorer outcome. Linear mixed-effects models were conducted with test score as the outcome and carrier status, age and gender as fixed effects and family as a random effect. Cohen’s d represents the standardised difference in trait score between ND-CNV carriers and controls adjusted for age and gender, scores were categorised into effect size descriptor categories; 0.00-0.19 negligible, 0.20-0.49 small, 0.50-0.79 medium, 0.80+ large. # denotes composite scores. *Survives Benjamini-Hochberg false discovery rate 0.05 correction

| Trait | ND-CNV carriers |  |  |  | Controls |  |  |  | Group difference |  |  |  |
| --- | --- | --- | --- | --- | --- | --- | --- | --- | --- | --- | --- | --- |
|  | N | z | 95% CI | SD | N | z | 95% CI | SD | p | d | 95% CI | Descriptor |
| **COGNITION** |  |  |  |  |  |  |  |  |  |  |  |  |
| FSIQ# | 237 | -1.48 | -1.64,-1.33 | 1.20 | 100 | 0.00 | -0.20,0.20 | 1.00 | $<1.00 \times 10^{-15*}$ | 1.30 | 1.04,1.56 | Large |
| PIQ# | 238 | -1.05 | -1.18,-0.91 | 1.05 | 99 | 0.00 | -0.20,0.20 | 1.00 | $<1.00 \times 10^{-15*}$ | 1.01 | 0.76,1.26 | Large |
| VIQ# | 239 | -1.43 | -1.58,-1.28 | 1.18 | 100 | 0.00 | -0.20,0.20 | 1.00 | $<1.00 \times 10^{-15*}$ | 1.26 | 1.00,1.52 | Large |
| Non-verbal reasoning | 238 | -1.01 | -1.16,-0.87 | 1.14 | 100 | 0.00 | -0.20,0.20 | 1.00 | $<1.00 \times 10^{-15*}$ | 0.93 | 0.68,1.17 | Large |
| Perceptual organisation | 237 | -0.84 | -0.96,-0.71 | 0.98 | 99 | 0.00 | -0.20,0.20 | 1.00 | $3.49 \times 10^{-14*}$ | 0.85 | 0.60,1.10 | Large |
| Verbal knowledge | 237 | -1.24 | -1.38,-1.11 | 1.04 | 100 | 0.00 | -0.20,0.20 | 1.00 | $<1.00 \times 10^{-15*}$ | 1.22 | 0.96,1.47 | Large |
| Verbal reasoning | 238 | -1.21 | -1.37,-1.06 | 1.20 | 100 | 0.00 | -0.20,0.20 | 1.00 | $<1.00 \times 10^{-15*}$ | 1.07 | 0.81,1.32 | Large |
| Set-shifting | 198 | -0.72 | -0.89,-0.56 | 1.14 | 87 | 0.00 | -0.21,0.21 | 1.00 | $5.99 \times 10^{-10*}$ | 0.66 | 0.40,0.92 | Medium |
| Spatial working memory | 212 | -0.81 | -0.96,-0.66 | 1.09 | 64 | 0.00 | -0.25,0.25 | 1.00 | $6.50 \times 10^{-08*}$ | 0.76 | 0.47,1.05 | Medium |
| Spatial planning | 170 | -0.58 | -0.72,-0.44 | 0.94 | 60 | 0.00 | -0.26,0.26 | 1.00 | $4.08 \times 10^{-05*}$ | 0.61 | 0.31,0.91 | Medium |
| Sustained attention | 155 | -1.01 | -1.21,-0.80 | 1.27 | 57 | 0.00 | -0.27,0.27 | 1.00 | $4.55 \times 10^{-08*}$ | 0.84 | 0.52,1.16 | Large |
| Processing speed | 194 | -0.39 | -0.54,-0.24 | 1.09 | 62 | 0.00 | -0.25,0.25 | 1.00 | $8.95 \times 10^{-03*}$ | 0.37 | 0.08,0.66 | Small |
| **PSYCHIATRIC TRAITS** |  |  |  |  |  |  |  |  |  |  |  |  |
| Total CAPA symptom count# | 233 | -1.23 | -1.38,-1.08 | 1.18 | 75 | 0.00 | -0.23,0.23 | 1.00 | $<1.00 \times 10^{-15*}$ | 1.09 | 0.81,1.36 | Large |
| Anxiety CAPA subscale | 233 | -0.85 | -1.04,-0.67 | 1.45 | 75 | 0.00 | -0.23,0.23 | 1.00 | $1.00 \times 10^{-06*}$ | 0.63 | 0.37,0.90 | Medium |
| ADHD CAPA subscale | 233 | -1.20 | -1.36,-1.04 | 1.23 | 75 | 0.00 | -0.23,0.23 | 1.00 | $1.82 \times 10^{-14*}$ | 1.02 | 0.74,1.30 | Large |
| Mood CAPA subscale | 233 | -0.60 | -0.74,-0.45 | 1.14 | 75 | 0.00 | -0.23,0.23 | 1.00 | $4.43 \times 10^{-5*}$ | 0.54 | 0.27,0.80 | Medium |
| OCD CAPA subscale | 233 | -0.67 | -0.92,-0.43 | 1.88 | 75 | 0.00 | -0.23,0.23 | 1.00 | $1.30 \times 10^{-3*}$ | 0.40 | 0.13,0.66 | Small |
| ODD CAPA subscale | 233 | -0.71 | -0.87,-0.56 | 1.17 | 75 | 0.00 | -0.23,0.23 | 1.00 | $4.36 \times 10^{-6*}$ | 0.63 | 0.37,0.90 | Medium |
| Sleep CAPA subscale | 233 | -0.69 | -0.86,-0.52 | 1.33 | 75 | 0.00 | -0.23,0.23 | 1.00 | $3.45 \times 10^{-5*}$ | 0.55 | 0.28,0.81 | Medium |
| Subclinical psychotic experiences (child CAPA) | 244 | -0.55 | -0.84,-0.27 | 2.27 | 83 | 0.00 | -0.22,0.22 | 1.00 | $2.97 \times 10^{-2*}$ | 0.27 | 0.02,0.52 | Small |
| **FUNCTIONING** |  |  |  |  |  |  |  |  |  |  |  |  |
| General functioning | 138 | -1.36 | -1.52,-1.20 | 0.96 | 56 | 0.00 | -0.27,0.27 | 1.00 | $<1.00 \times 10^{-15*}$ | 1.40 | 1.05,1.75 | Large |
| Social functioning | 128 | -1.60 | -1.77,-1.42 | 1.01 | 56 | 0.00 | -0.27,0.27 | 1.00 | $<1.00 \times 10^{-15*}$ | 1.60 | 1.22,1.96 | Large |
| **PSYCHOPATHOLOGY** |  |  |  |  |  |  |  |  |  |  |  |  |
| ASD traits | 227 | -2.21 | -2.40,-2.03 | 1.39 | 86 | 0.00 | -0.21,0.21 | 1.00 | $<1.00 \times 10^{-15*}$ | 1.71 | 1.42,2.01 | Large |
| Motor coordination | 226 | -1.89 | -2.05,-1.73 | 1.23 | 86 | 0.00 | -0.21,0.21 | 1.00 | $<1.00 \times 10^{-15*}$ | 1.62 | 1.33,1.91 | Large |
| SDQ total (caregiver report)# | 229 | -1.59 | -1.72,-1.45 | 1.05 | 88 | 0.00 | -0.21,0.21 | 1.00 | $<1.00 \times 10^{-15*}$ | 1.54 | 1.25,1.82 | Large |
| Conduct SDQ subscale (caregiver report) | 229 | -0.60 | -0.74,-0.46 | 1.09 | 88 | 0.00 | -0.21,0.21 | 1.00 | $4.97 \times 10^{-6*}$ | 0.57 | 0.32,0.82 | Medium |
| Emotional SDQ subscale (caregiver report) | 229 | -0.79 | -0.93,-0.65 | 1.06 | 88 | 0.00 | -0.21,0.21 | 1.00 | $7.83 \times 10^{-10*}$ | 0.76 | 0.50,1.01 | Medium |
| Hyperactivity SDQ subscale (caregiver report) | 229 | -2.27 | -2.45,-2.09 | 1.39 | 88 | 0.00 | -0.21,0.21 | 1.00 | $<1.00 \times 10^{-15*}$ | 1.76 | 1.46,2.05 | Large |
| Peer SDQ subscale (caregiver report) | 229 | -1.39 | -1.54,-1.24 | 1.14 | 88 | 0.00 | -0.21,0.21 | 1.00 | $<1.00 \times 10^{-15*}$ | 1.27 | 0.99,1.54 | Large |
| Prosocial SDQ subscale (caregiver report) | 229 | -0.87 | -1.02,-0.73 | 1.09 | 88 | 0.00 | -0.21,0.21 | 1.00 | $1.56 \times 10^{-11*}$ | 0.82 | 0.56,1.08 | Large |
| SDQ total (teacher report)# | 138 | -1.06 | -1.23,-0.90 | 0.99 | 52 | 0.00 | -0.28,0.28 | 1.00 | $1.36 \times 10^{-11*}$ | 1.08 | 0.73,1.42 | Large |
| Conduct SDQ subscale (teacher report) | 138 | -0.45 | -0.65,-0.25 | 1.17 | 52 | 0.00 | -0.28,0.28 | 1.00 | $1.31 \times 10^{-3*}$ | 0.40 | 0.08,0.72 | Small |
| Emotional SDQ subscale (teacher report) | 138 | -0.53 | -0.69,-0.37 | 0.97 | 52 | 0.00 | -0.28,0.28 | 1.00 | $7.54 \times 10^{-4*}$ | 0.54 | 0.22,0.87 | Medium |
| Hyperactivity SDQ subscale (teacher report) | 138 | -1.36 | -1.61,-1.12 | 1.45 | 52 | 0.00 | -0.28,0.28 | 1.00 | $1.84 \times 10^{-10*}$ | 1.02 | 0.67,1.36 | Large |
| Peer SDQ subscale (teacher report) | 138 | -0.86 | -1.06,-0.66 | 1.18 | 52 | 0.00 | -0.28,0.28 | 1.00 | $1.96 \times 10^{-6*}$ | 0.76 | 0.43,1.09 | Medium |
| Prosocial SDQ subscale<br>(teacher report) | 138 | -0.96 | -1.19,-0.73 | 1.3<br>7 | 52 | 0.00 | -0.28,0.28 | 1.00 | $2.30 \times 10^{-6*}$ | 0.7<br>6 | 0.42,1.09 | Medium |

**Figure 2:**
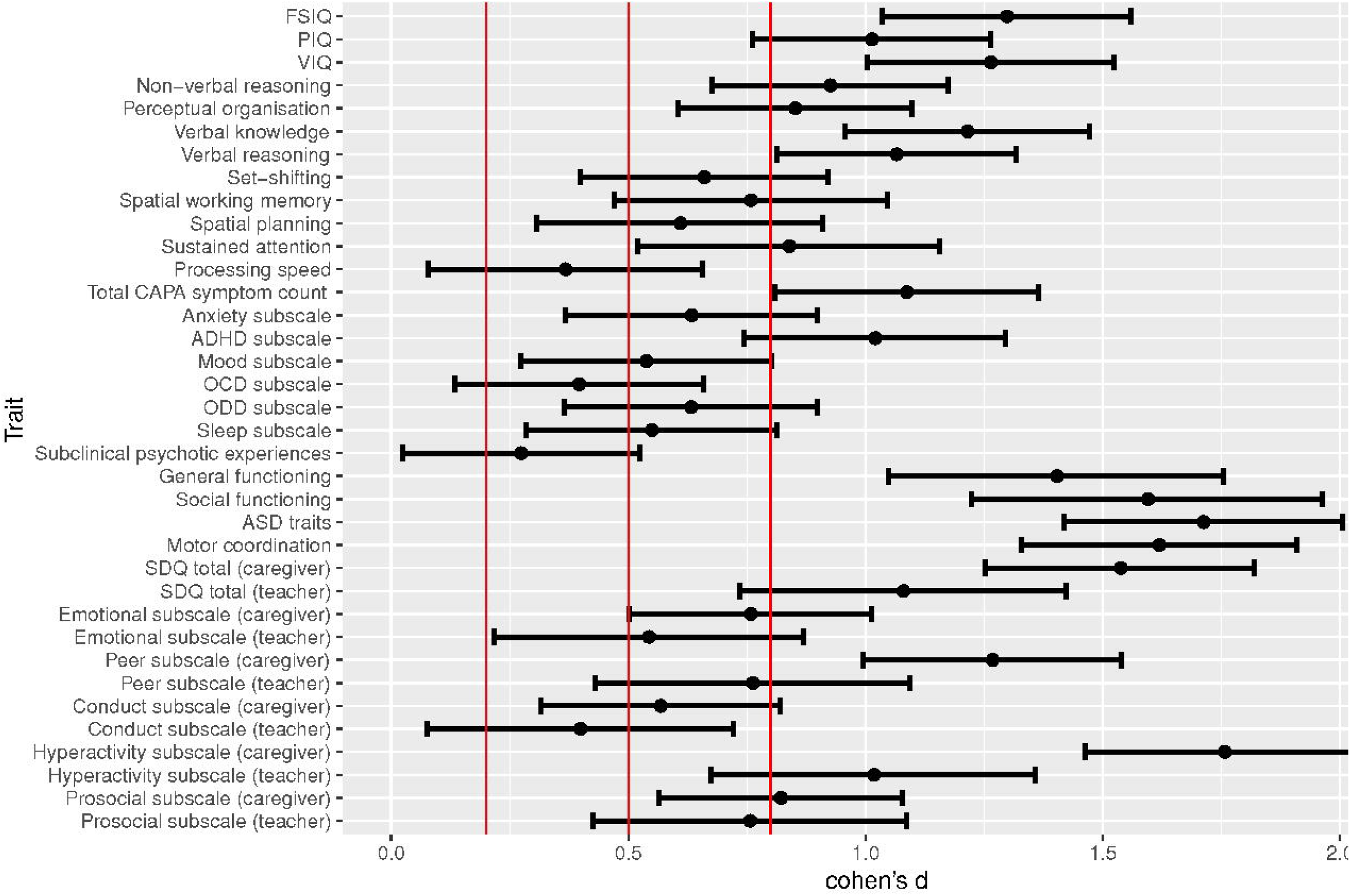
Standardised difference between ND-CNV carriers and controls on quantitative cognitive and behavioural traits. FSIQ, Full Scale Intelligence Quotient; PIQ, Performance Intelligence Quotient; VIQ, Verbal Intelligence Quotient; CAPA, Child and Adolescent Psychiatric Assessment; ADHD, Attention Deficit Hyperactivity Disorder; OCD, Obsessive Compulsive Disorder; ODD, Oppositional Defiant Disorder; ASD, Autism Spectrum Disorder; SDQ, Strengths and Difficulties Questionnaire. Cohen’s d represents the standardised difference in trait between ND-CNV carriers and controls adjusted for age and gender. The red lines denote effect size descriptor categories^51^; 0.00-0.19 negligible, 0.20-0.49 small, 0.50-0.79 medium, 0.80+ large.

*Aim 2:* Within ND-CNV carriers, mean performance (adjusted for age and gender) on phenotypic traits for each ND-CNV are visualised in Figure 3, where distinct profiles are apparent. Regarding phenotypic associations, 2 clusters can be distinguished; *neurodevelopmental traits* (Figure 3, Box B); and *mental health and cognitive comorbidities* (Figure 3, Box A). All genotypes showed evidence of strong impairments in the *neurodevelopmental traits* cluster, whereas level of impairment within the *mental health and cognitive comorbidities* cluster was less and more variable across ND-CNVs (the traits that comprise each cluster are shown in Figure 3, and see the Supplementary Table 1 for z-scores for each ND-CNV). The dendrogram on Figure 3 shows the pattern of ND-CNV clustering, there was no strong evidence that deletion variants differed in profile from duplication variants, or that deletions and duplications at the same loci differed in profile.

**Figure 3:**
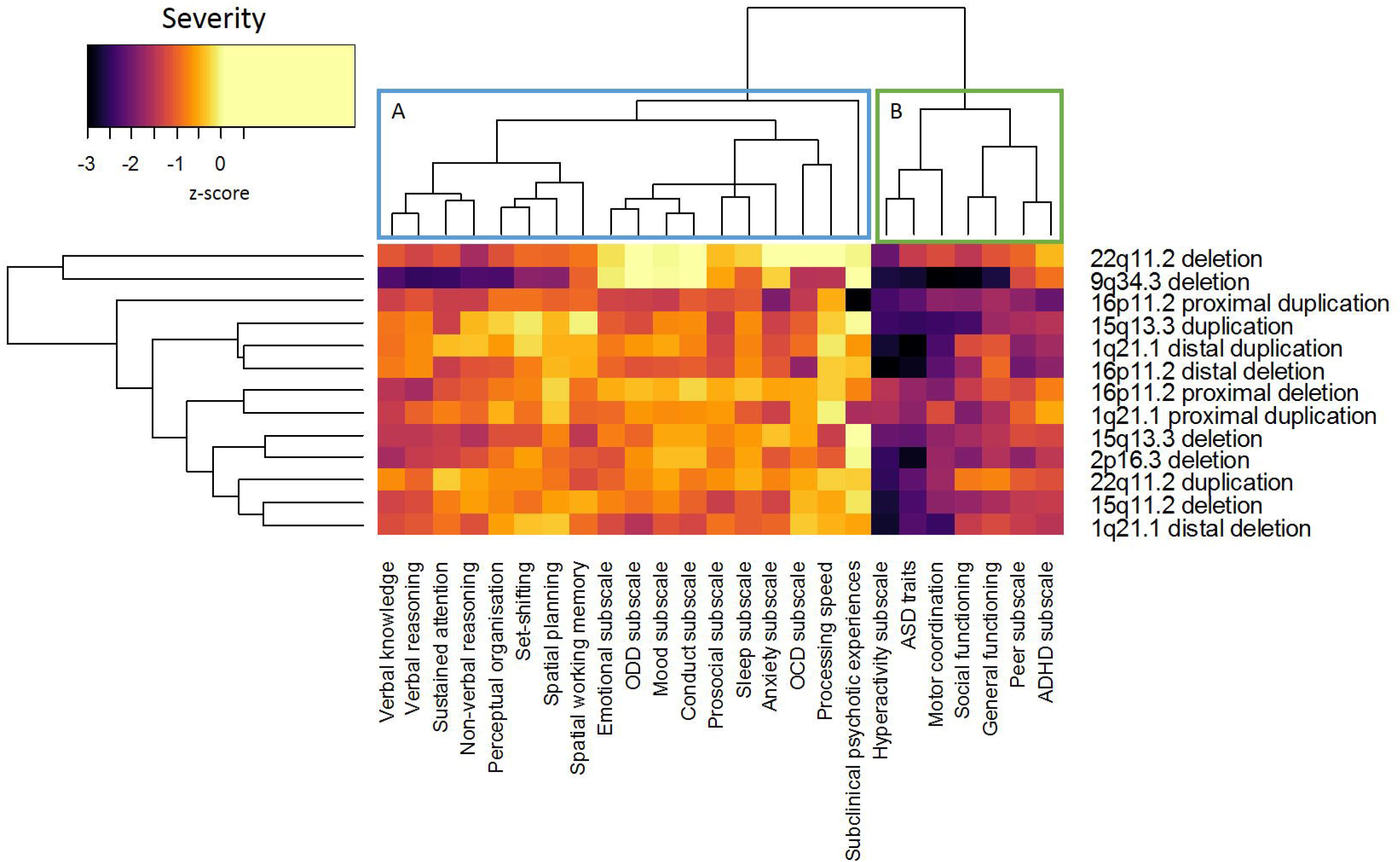
Phenotypic profiles of individual ND-CNV genotypes. ADHD, Attention Deficit Hyperactivity Disorder; OCD, Obsessive Compulsive Disorder; ODD, Oppositional Defiant Disorder; ASD, Autism Spectrum Disorder. Test scores were transformed so that their distribution approximated the normal distribution as closely as possible. Transformed scores were standardised into z scores using the means and SDs of the control group as reference and adjusted for age and gender, and were constructed so that a negative score denoted a poorer outcome. Cream colour represents a z score difference of zero between the ND-CNV group and controls, whereas yellow through to orange, red purple and black represents a deficit in the CNV group relative to controls. Hierarchical clustering, for the purposes of presentation (indicated by the dendrogram), was performed using Ward’s method and Euclidian distance. Domains clustered into two groups; mental health and cognitive comorbidities (cluster A, blue box) and neurodevelopmental traits (cluster B, green box)

In terms of the overall phenotypic profile there was evidence of both qualitative and quantitative differences between genotypes. Both tests of significance for rank concordance and discordance were significant for both analyses of qualitative (Friedman chi-squared = 177.39, p<1.00×10^−15^; Kendall F=15.81, p<1.00×10^−15^) and quantitative effects (Friedman chi-squared = 53.04, p=4.06×10^−7^; Kendall F=5.15, p<8.72×10^−8^). These findings indicate that, although significant quantitative and qualitative differences exist, the converse is true in that qualitative and quantitative similarities also exist. We therefore conclude effects for both qualitative as well as quantitative differences between genotypes are moderate, and overall our data supports Model 4 (Figure 1). A sensitivity analysis was conducted excluding individuals with 9q34.3 deletion or 22q11.2 deletion as the dendrogram (Figure 3) indicated their phenotypic profiles stood apart from the other ND-CNVs and that this could drive the differences we found. However, excluding these two groups did not change our finding of moderate qualitative and a quantitative differences between genotypes, and did not change the hierarchical clustering of traits into *neurodevelopmental traits* and *mental health and cognitive comorbidities*. We conducted further sensitivity analysis to confirm that our findings were not driven by a) by one of the two clusters specifically or b) overlap between different phenotypic measures (Supplementary Materials). We found that a) moderate qualitative and quantitative differences existed within both the *neurodevelopmental traits* as well as the *mental health and cognitive comorbidities* clusters, indicating that both phenotypic clusters drive qualitative and quantitative differences; b) findings remained the same when analyses were conducted on a second set of rankings based on sub-cluster scores, therefore taking account of overlapping correlated phenotypic traits. Sub-clusters were identified from the hierarchical clustering conducted for Figure 3 (listed in Supplementary Materials).

At the level of individual phenotypic traits quantitative differences were found with genotype predicting between 5-20% of variance (Eta-squared effect size) in impairment within ND-CNV carriers depending on the specific trait (Table 4). The effect of genotype significantly predicted impairment severity in some traits; FSIQ, PIQ, VIQ, all the IQ subtests, spatial planning, processing speed, total CAPA symptom count including the anxiety, ADHD and ODD subscales, subclinical psychotic experiences, social functioning, ASD traits, motor coordination, SDQ total including conduct, hyperactivity and prosocial subscales (Table 4). However, for set-shifting ability, spatial working memory, sustained attention, mood CAPA subscale, OCD CAPA subscale, sleep CAPA subscale, general functioning, emotional SDQ subscale, and the peer SDQ subscale, the effect of genotype was not significant. For these analyses, p-values equal or less than 0.02 survived BH-FDR 0.05 correction. These findings remained largely the same when we included family ethnic background and family income in the analysis, with genotype explaining 5-21% of variation in phenotypic outcome, depending on the trait (see Supplementary Table 3).

**Table 4:**
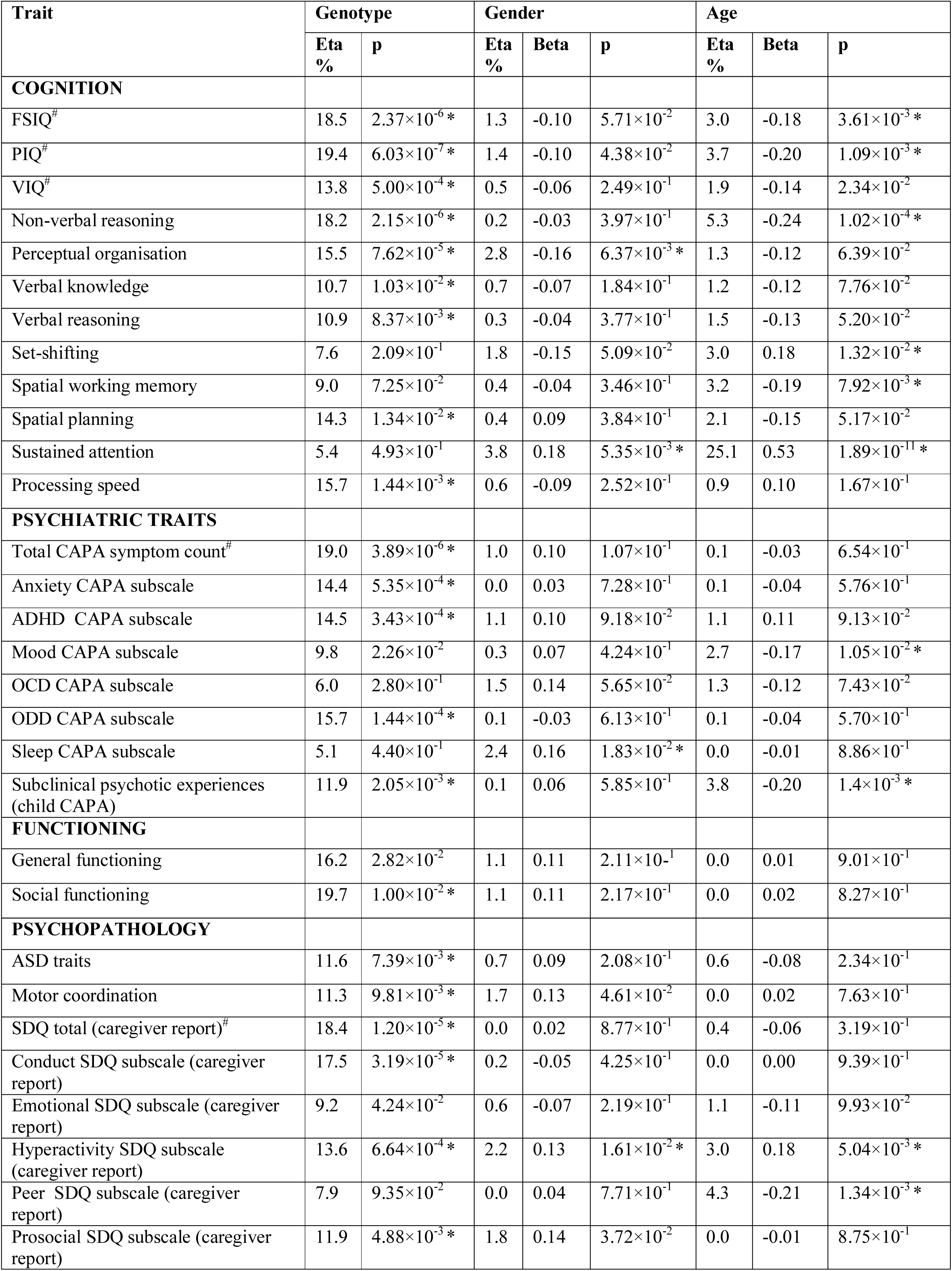
Effect size of genotype, age, and gender on phenotypic outcomes. Eta, eta-squared, FSIQ, Full Scale Intelligence Quotient; PIQ, Performance Intelligence Quotient; VIQ, Verbal Intelligence Quotient; CAPA, Child and Adolescent Psychiatric Assessment; ADHD, Attention Deficit Hyperactivity Disorder; OCD, Obsessive Compulsive Disorder; ODD, Oppositional Defiant Disorder; ASD, Autism Spectrum Disorder; SDQ, Strengths and Difficulties Questionnaire. P-values, eta-squared values were derived from ANCOVA analyses examining the effect of genotype, age and gender. Standardised beta values were derived from linear regression models. For gender a positive beta value indicated that males had a higher score compared to females, for age a positive beta value indicated that the score increased with age. # denotes composite scores. *Survives Benjamini-Hochberg false discovery rate 0.05 correction

*Aim 3:* The phenotypic profile of ND-CNV carriers was influenced by age (accounting for 0-25% of variance depending on trait) with deficits in some traits becoming reduced in older children: the hyperactivity SDQ subscale (β = 0.18, p=5.04×10^−3^), sustained attention (β = 0.53, p=1.89×10^−11^ and higher executive function (set-shifting, β = 0.18, p=1.32×10^−2^). Deficits in other traits were found to be greater in older children; FSIQ (β = −0.18, p=3.61×10^−3^), spatial working memory (β = −0.10, p=7.92×10^−3^), mood CAPA subscale (β = −0.17, p=1.05×10^−2^), subclinical psychotic experiences (β = −0.20, p=1.40×10^−3^), and the peer subscale of the SDQ (β = −0.21, p=1.34×10^−3^). We conducted interaction analysis to evaluate, for the traits we found to be age related, whether there was evidence of differences in the relationship between age and phenotypic outcome between ND-CNV carriers and controls. An interaction between age and group status (ND-CNV carriers vs controls) was found only for the CAPA mood subscale score (p=4.38×10^−2^), indicating that with increasing age mood problems develop at a greater rate in ND-CNV carriers relative to controls. For the other traits (hyperactivity, spatial working memory, sustained attention, set-shifting, FSIQ, spatial working memory, subclinical psychotic experiences and the peer SDQ subscale) no evidence of differential development with age was found between ND-CNV carriers and controls. Gender was found to influence phenotypic outcomes in ND-CNV carriers, but accounted for little variation (0-4% depending on trait). Males had greater deficits on the hyperactivity SDQ subscale (β = 0.17, p=1.61×10^−2^), sleep CAPA subscale (β = 0.16, p=1.83×10^−2^) and sustained attention (β = 0.18, p=5.35×10^−3^) than females, but they performed better on the PIQ perceptual organisation subtest (β = −0.16, p=6.37×10^−3^). For these analyses, p-values equal or less than 0.02 survived BH-FDR 0.05 correction. These results remained largely the same when taking into account family ethnicity and family income (see Supplementary Table 3).

## Discussion

The IMAGINE-ID cohort allowed us to conduct one of the largest studies to define genotype-phenotype relationships across a range of ND-CNV loci. Overall our findings support a model whereby these ND-CNVs have a broadly general effect on phenotypic outcome, but specific effects can be identified, albeit accounting for a low proportion of variance (5-20% depending on trait). Some traits had similar levels of impairment across all genotypes (e.g. mood problems, sleep impairments, peer problems, and sustained attention) whereas for other traits there was more evidence of genotype specific patterns (e.g., IQ, spatial planning, processing speed, subclinical psychotic experiences, ASD traits, motor coordination total psychiatric symptomatology, particularly anxiety, ADHD, and conduct related traits). Phenotypic differences between ND-CNVs were found to be both quantitative and as well as qualitative in nature (Model 4, Figure 1). Hierarchical cluster analysis of phenotypic traits identified two clusters; *neurodevelopmental traits* that were strongly impaired across CNVs, and *mental health and cognitive comorbidities* where impairment was generally less and more variable across the genotypes. ND-CNVs affect biological pathways that impact risk of developmental impairment and this impairment differs in magnitude by genotype, but the unique gene content of each ND-CNV also appears to mould the specific psychiatric, cognitive and other manifestations. As a group, children with a ND-CNV were found to be at very high risk of developing psychiatric disorder, with 79.8% having at least one psychiatric diagnosis. Moreover, using a broad multi-informant approach we found that ND-CNV carriers were impaired relative to their siblings across all the psychiatric, neurodevelopmental, psychopathological, cognitive, social, sleep and motor domains assessed. This patient group clearly warrants clinical and educational attention and intervention.

ND-CNV carriers were found to be at increased risk for a range of psychiatric disorders (OR=13.8 for any disorder), including ADHD (OR=6.9), anxiety disorder (OR=2.9), ASD (OR=44.1), ODD (OR=43.6), and tic disorders (OR could not be calculated as no controls were affected). All the ND-CNV carriers were impaired across all behavioural and cognitive traits measured, the strongest trait differences found between ND-CNV carriers and controls included ASD symptom count (d=1.71), hyperactivity (d=1.76), social functioning (d=1.60) and motor coordination (d=1.62). Motor coordination is a domain that has been relatively understudied in the context of ND-CNV carriers, but recent studies indicate that it is an antecedent^35^ of, and indexes, psychiatric disorder^36^ in ND-CNV carriers. Our teacher-report measures confirmed that neuropsychiatric impairments were present in multiple settings, indicating pervasiveness. Our findings of broad ranging impairments is consistent with studies of common polygenic risk^37^ and familial risk^38,39^ of psychiatric disorder, that find that genetic risk is associated with disrupted childhood neurodevelopment across several domains. Psychiatric disorder was present in control siblings at rates in line with previous population studies that have used the same instruments as in the current study ^40,41^ as well as with previous studies that have compared ND-CNV carriers to controls^42,43^. Some previous studies of 22q11.2 deletion carriers have contrasted to community controls, but the effect sizes we find when we contrast to sibling controls are broadly similar^44,45^.

Strikingly, the specific effect of CNV genotype only accounted for 5-20% of variation in outcome depending on phenotypic trait, indicating that the majority of variance is explained by additional factors. We found that age was a predictor of outcome for several traits, both ADHD symptoms and deficits in our cognitive measures of sustained attention and executive function decreased with age, whereas IQ deficits, spatial working memory, mood symptoms, subclinical psychotic experiences and peer problems increased with age. These trends are in line with general population studies^40,46^. We do find that increase of mood symptoms with age was greater in ND-CNV carriers relative to controls. The other phenotypic traits however, although impaired in ND-CNV carriers, showed comparable trends with increasing age as in in the controls. These initial cross-sectional findings illustrate the importance of having a control group when viewing genotype-phenotype relationships through a developmental lens^17,47^. These cross-sectional findings warrant future longitudinal studies. Although we found gender differences in neurodevelopmental traits in line with findings within the general population^48^, the proportion of variance explained was low (<5%). This may reflect that male-to-female ratios for conditions such as autism are reduced in populations with intellectual disability^49^. Further research will be required to understand what genetic and environmental factors underlie the remaining, unexplained variation, in outcome.

The high prevalence of psychiatric disorders and the finding that ND-CNV carriers were impaired across all the cognitive, motor and psychopathological measures assessed highlight that children with ND-CNVs require coordinated multidisciplinary care to address a range of psychiatric, psychological, motor coordination, sleep and social and educational needs. This warrants a step change in current clinical service provision, and calls for greater awareness of this new patient group amongst clinicians and educators. The commonalities we are finding in clinical outcomes across ND-CNVs suggest this group could benefit from the development of a dedicated clinical care pathway, which would provide psychoeducation about the broad range of associated risk alongside tailored monitoring of more genotype-specific vulnerabilities. Support and intervention plans for children with a ND-CNV need to consider the child’s behaviour in educational and peer contexts as well as address behaviour exhibited in the home or clinic. The presence of commonalities in clinical outcomes also indicates that genomic risk for neurodevelopmental conditions impacts shared biological processes which could be targeted for pharmacological intervention. In this light, it is noteworthy that several ND-CNVs have been linked to synaptic dysfunction^16^. The broad ranging phenotypic outcomes associated with ND-CNVs indicates that genotype-phenotype relationships have a complex architecture. Current research efforts that use genetic first approaches in human studies and animal models as a way of identifying direct causal pathways from genotype to psychiatric disorder via intermediary phenotypes need to take account of these complexities, the use of an endophenotype approach should be cautioned^50^. Systems biology and network approaches are needed to globally capture the architecture of genotype-phenotype relationships. Efforts focusing on single causal pathways are likely to only provide limited research and clinical benefit.

## Limitations

Individuals had to have a known genetic diagnosis to take part, the study would therefore not capture asymptomatic individuals who carry ND-CNVs. Therefore, the true phenotype in the general population is likely to be less severe than what we report, with bias being greater for those ND-CNVs with a lower penetrance. However, it is important to put our study in context with wider research. Large population-based studies have examined the phenotype of ND-CNVs in adults from the general population^7,9^, but the present study gives unique insights into the development of children at the more severe end of the phenotypic spectrum who are most likely to engage with health services and be in need of clinical and educational support. Some of our findings could reflect ascertainment bias, in that carriers with severe developmental delay are more likely to be referred to medical genetic clinics for testing. However, we found that the differences between ND-CNV carriers and controls remained significant after controlling for IQ. Due to our sample size we may be underpowered to detect more subtle genotype-phenotype relationships, however this is unlikely to affect our main conclusions that ND-CNVs have large pleiotropic effects on childhood development and that although specific genotype-phenotype relationships exist within ND-CNV carriers the effect size is relatively low. IMAGINE-ID is a nationwide study, however, to increase power in future studies, multinational collaborations will be needed. Initiatives such as the pan-European “Maximising Impact of research in NeuroDevelopmental DisorderS” MINDDS network (https://mindds.eu/) provide a springboard for developing international studies of ND-CNV carriers.

## Conclusion

Our findings provide evidence of specific genotype-phenotype relationships within CNV carriers both in terms of quantitative and qualitative differences. However, although differences can be identified, these account for a low proportion of variance and therefore we conclude that different genotypes do not result in discrete forms of neurodevelopmental disorders. Our dimensional approach facilitated the investigation of genotype-phenotype relationships beyond categorical psychiatric diagnosis. Using a multi-informant deep phenotyping approach we found that genomic risk for psychiatric disorder had wide ranging effects on childhood development spanning a range of cognitive and behavioural domains. Our findings highlight that there are core neurodevelopmental traits that are strongly impaired across all ND-CNV carriers, but additionally ND-CNV carriers are also affected by broad-ranging mental health and cognitive comorbidities. This suggests that multiple processes and neural circuits are affected by ND-CNVS. Future research into the relationship between genotype and psychiatric outcomes via intermediary endophenotypes needs to consider this when interpreting findings. Early detection of children with ND-CNVs is warranted to a) investigate antecedents and developmental course of neuropsychiatric impairments, b) add to the understanding of how genomic risk manifests, c) inform early intervention programs.

## Acknowledgements

We would like to thank all the children, families and teachers who took part in the IMAGINE-ID study, as well as all the support we have had from NHS medical genetic clinics, and support charities including Unique and Max Appeal. We thank all members of the IMAGINE-ID consortium for their contributions. We thank Matthew Sopp, Alice Robinson, Sinéad Ray, Nicola Lewis, Sarah Law, Sophie Andrews, Aimée Challenger, Poppy Sloane, Alice Walsh, Keziah Fish, Amy Ilsley, Stephen Naughton, Rachel Tompkins, Ciara Walker, Nadia Pantouw, Samantha Bowen, Hannah Pendlebury, Chloe Sheldon, Emily Green, Umaya Prasad, Joshua Roberts, Jessica Townsend and Beth Hughes for their roles in data collection. We thank Nigel Williams, Andrew Cuthbert and Alexandra Evans for their support in interpreting genotype information, Hayley Moss for research management support and Karen Bradley for administrative support. We thank the core laboratory team of the Division of Psychological Medicine and Clinical Neurosciences laboratory for DNA sample management and genotyping. We also thank the National Centre for Mental Health, a collaboration between Cardiff, Swansea and Bangor Universities, for their support.

This work was funded by Medical Research Council grants MR/L011166/1 and MR/N022572/1, and by the Waterloo Foundation.

## Contributors

SJRAC, MvdB, JH and MO drafted the manuscript, which was reviewed and approved by all other authors. MvdB and SJRAC coordinated the data collection. SJRAC, MvdB, PH, JH and MO were involved in the analysis and interpretation of the data. The IMAGINE-ID study was designed by MvdB and JH (deep phenotyping study) and DS and LR (online study).

