## Supplemental Information for "Genotype-phenotype relationships in children with Copy Number Variants associated with high neuropsychiatric risk: Findings from the case-control IMAGINE-ID cohort in the United Kingdom"

#### Full assessment details

*Cognition*

IQ was assessed using the Wechsler Abbreviated Scale of Intelligence (WASI)[1], from which Full Scale IQ (FSIQ), Verbal IQ (VIQ) and Performance IQ (PIQ) were derived. To examine specific cognitive traits, subtest performance scores were derived which captured verbal knowledge (vocabulary subtest), verbal reasoning (similarities subtest), perceptual organisation (block design subtest) and non-verbal reasoning (matrix reasoning subtest).

Further specific cognitive traits were assessed using the Wisconsin Card Sorting Test (WSCT)[2] where number of perseverative errors measured the set-shifting ability aspect of executive function, as well as the following CANTAB (Cambridge Neuropsychological Test Automated Battery)[3] tests: spatial working memory (SWM) an executive function task; stockings of Cambridge (SOC) which measures spatial planning and is an executive function task; five choice reaction time (RTI) which measures processing speed; match-to-sample task (MTS) a test of visual attention; and rapid visual information processing (RVP) a measure of sustained attention.

*Psychopathology*

Psychiatric symptomatology, during the last 3 months was assessed using the Child and Adolescent Psychiatric Assessment (CAPA)[4] by means of semi-structured interview with the primary carer (parent CAPA). Psychosis spectrum outcomes, including subclinical psychotic experiences and psychotic disorder, were assessed using the psychosis section of the child report CAPA. Criteria were applied to establish DSM-IV-TR [5] diagnosis, and symptom subscale scores were also derived. All interviews were audio recorded and were consensus coded by two psychologists who received supervision from a child and adolescent psychiatrist. The CAPA was chosen as it has been used successfully in assessing psychiatric disorder in UK cohorts of children with neurodevelopmental conditions including ASD[6] and 22q11.2 deletion syndrome[7,8] .

Autism Spectrum Disorder (ASD) traits were assessed using the Social Communication Questionnaire (SCQ) [9], which was completed by the primary carer. Total scores can range from 0 to 39. In our analysis we used the recommended [10] score of 15 or higher as indicating probable ASD.

Individuals were classified as having any psychiatric disorder if they met criteria on either the SCQ or CAPA.

Motor coordination impairment was assessed using the Developmental Coordination Disorder Questionnaire (DCDQ)[11], which was completed by the primary carer.

A dimensional measure of broader child psychopathology was assessed using the Strengths and Difficulties Questionnaire (SDQ)[12], which comprises conduct, emotional, hyperactivity, peer and prosocial subscales. Conduct, emotional, hyperactivity and peer subscales were combined into a total score. The prosocial scale is a positive scale, and does not contribute to the total score. The SDQ was completed by the primary carer and as well as the child's teacher. Teacher report data was gathered after the home visit, caregivers had to consent for the school to be contacted, and following this teachers completed a consent form and postal questionnaire.

*Functioning and educational outcomes*

General functioning, Children's Global Assessment Scale (CGAS)[13], and social functioning, Social and Occupational Functioning Assessment Scale (SOFAS), were conducted by two psychologists reaching consensus on functioning of the child as observed during the home visit. These measures were added after an initial pilot phase of the study, and are therefore not available for all individuals.

*Summary table*

| Trait | Measure | Informant | **Composite score** |
|---|---|---|---|
| FSIQ | WASI | Child | Yes |
| PIQ | WASI | Child | Yes |
| VIQ | WASI | Child | Yes |
| Non-verbal reasoning | WASI Matrix Reasoning subtest | Child | No |
| Perceptual organisation | WASI Block Design subtest | Child | No |
| Verbal knowledge | WASI Vocabulalry subtest | Child | No |
| Verbal reasoning | WASI Similarities subtest | Child | No |
| Set-shifting | WCST perservative errors | Child | No |
| Spatial working memory | CANTAB Spatial Working Memory subtest | Child | No |
| Spatial planning | CANTAB Stockings of Cambridge subtest | Child | No |
| Sustained attention | CANTAB Rapid Visual Processing subtest | Child | No |
| Processing speed | CANTAB 5 Choice Reaction Time | Child | No |
| Total CAPA symptom count | CAPA | Carer | Yes |
| Anxiety subscale | CAPA | Carer | No |
| ADHD subscale | CAPA | Carer | No |
| Mood subscale | CAPA | Carer | No |
| OCD subscale | CAPA | Carer | No |
| ODD subscale | CAPA | Carer | No |
| Sleep subscale | CAPA | Carer | No |
| Subclinical psychotic experiences | CAPA | Child | No |
| General functioning | C-GAS | Research psychologist | No |
| Social functioning | SOFAS | Research psychologist | No |
| ASD traits | SCQ | Carer | No |
| Motor coordination | DCDQ | Carer | No |
| SDQ total | SDQ | Carer and teacher | Yes |
| Conduct subscale | SDQ | Carer and teacher | No |
| Emotional subscale | SDQ | Carer and teacher | No |
| Hyperactivity subscale | SDQ | Carer and teacher | No |
| Peer subscale | SDQ | Carer and teacher | No |
| Prosocial subscale | SDQ | Carer and teacher | No |

**Qualitative and quantitative sensitivity analyses**

We wanted to confirm that our findings were not driven by just one cluster, so we conducted analysis of qualitative and quantitative differences within both the *neurodevelopmental traits* and *mental health and cognitive comorbidities* clusters.

For the *neurodevelopmental traits* cluster, both tests of significance for rank concordance and discordance were significant for both analyses of qualitative (Friedman chi-squared = 48.13, $p=1.11\times10^{-8}$; Kendall F=19.34, $p=5.28\times10^{-13}$) and quantitative effects (Friedman chi-squared = 39.92, $p=7.42\times10^{-5}$; Kendall F=5.43, $p=2.28\times10^{-6}$).

For the *mental health and cognitive comorbidities* cluster, both tests of significance for rank concordance and discordance were significant for both analyses of qualitative (Friedman chi-squared = 53.84, $p=1.05\times10^{-5}$; Kendall F=3.87, $p=1.70\times10^{-6}$) and quantitative effects (Friedman chi-squared = 35.07, $p=4.57\times10^{-4}$; Kendall F=3.30, $p=2.36\times10^{-4}$).

Furthermore, we wanted to confirm that our findings were not driven by overlap between different phenotypic measures so we conducted rank based analyses on sub-cluster scores. Below are a list of sub-clusters which were derived from the hierarchical cluster analysis.

On a new set of rankings based on sub-cluster score, both tests of significance for rank concordance and discordance were significant for both analyses of qualitative (Friedman chi-squared = 57.48,

$p=1.45\times10^{-9}$; Kendall F=14.82, $p=1.06\times10^{-13}$) and quantitative effects (Friedman chi-squared = 24.35, $p=1.82\times10^{-2}$; Kendall F=2.33, $p=1.22\times10^{-2}$).

*Sub clusters*

Sub cluster 1: Verbal knowledge, verbal reasoning, sustained attention and non-verbal reasoning

Sub-cluster 2: Perceptual organisation, set-shifting, spatial planning and spatial working memory

Sub-cluster 3: Emotional subscale, ODD subscale, mood subscale and conduct subscale

Sub-cluster 4: Prosocial subscale and sleep subscale

Sub-cluster 5: Anxiety subscale

Sub-cluster 6: OCD subscale and processing speed

Sub-cluster 7: Subclinical psychotic experiences

Sub-cluster 8: Hyperactivity subscale, ASD traits and motor coordination

Sub-cluster 9: Social functioning, general functioning, peer subscale and ADHD subscale

#### Supplementary Tables

**Supplementary Table 1 Table:** Phenotypic trait z-scores and orginal scores for each ND-CNV group

FSIQ, Full Scale Intelligence Quotient; PIQ, Performance Intelligence Quotient; VIQ, Verbal Intelligence Quotient; CAPA, Child and Adolescent Psychiatric Assessment; ADHD, Attention Deficit Hyperactivity Disorder; OCD, Obsessive Compulsive Disorder; ODD, Oppositional Defiant Disorder; ASD, Autism Spectrum Disorder; SDQ, Strengths and Difficulties Questionnaire. Test scores were transformed so that their distribution approximated the normal distribution as closely as possible.Transformed scores were standardised into z scores using the means and SDs of the control group as reference and adjusted for age and gender, and were constructed so that a negative score denoted a poorer outcome. The original test scores are also presented. “Combined deletion group” brings together all the ND-CNV deletions, and “Combined duplication group” brings together all the ND-CNV duplications. The p-values indicate whether the z-score of the ND-CNV group differs from controls, and were derived from the linear mined models conducted in Aim 1 and are adjusted for multiple post hoc contrasts (Tukey’s method). P<0.05 are highlighted in green.Note contrasts for 9q34 deletion, 15q13.3 duplication, 16p11.2 distal deletion 1q21.1 proximal duplication and 2p16.3 deletion should be interpreted cautiously due to sample size, within this groups there was only power to detect large effect sized contrasts.

**Supplementary Table 2**: Quantitative cognitive and behavioural traits in controls and ND-CNV carriers with unconfirmed control siblings excluded

FSIQ, Full Scale Intelligence Quotient; PIQ, Performance Intelligence Quotient; VIQ, Verbal Intelligence Quotient; CAPA, Child and Adolescent Psychiatric Assessment; ADHD, Attention Deficit Hyperactivity Disorder; OCD, Obsessive Compulsive Disorder; ODD, Oppositional Defiant Disorder; ASD, Autism Spectrum Disorder; SDQ, Strengths and Difficulties Questionnaire. Test scores were transformed so that their distribution approximated the normal distribution as closely as possible. Transformed scores were standardised into z scores using the means and SDs of the control group as reference and adjusted for age and gender, and were constructed so that a negative score denoted a poorer outcome. Linear mixed-effects models were conducted with test score as the outcome and carrier status, age and gender as fixed effects and family as a random effect. Cohen’s d represents the standardised difference in trait score between ND-CNV carriers and controls adjusted for age and gender, scores were categorised into effect size descriptor categories; 0.00-0.19 negligible, 0.20-0.49 small, 0.50-0.79 medium, 0.80+ large.

*Survives Benjamini-Hochberg false discovery rate 0.05 correction

**Supplementary Table 3**: Effect size of genotype, age, and gender on phenotypic outcomes, controlled for ethnicity and family income.

Eta, eta-squared, FSIQ, Full Scale Intelligence Quotient; PIQ, Performance Intelligence Quotient; VIQ, Verbal Intelligence Quotient; CAPA, Child and Adolescent Psychiatric Assessment; ADHD, Attention Deficit Hyperactivity Disorder; OCD, Obsessive Compulsive Disorder; ODD, Oppositional Defiant Disorder; ASD, Autism Spectrum Disorder; SDQ, Strengths and Difficulties Questionnaire. P-values, eta-squared values were derived from ANCOVA analyses examining the effect of genotype, age and gender whilst controlling for ethnicity and family income. Standardised beta values were derived from linear regression models. For gender a positive beta value indicated that males had a higher score compared to females, for age a positive beta value indicated that the score increased with age.


1. Wechsler D. Manual for the Wechsler abbreviated intelligence scale (WASI); 1999.
2. Heaton R, Chelune G, Talley J, Kay G, Curtiss G. Wisconsin card sorting test manual revised and expanded. Lutz, FL: Psychological Assessment Resources. Inc; 1993.
3. CANTAB. CANTAB eclipse version 3. Cambridge: Cambridge Cognition; 2006.
4. Angold A, Prendergast M, Cox A, Harrington R, Simonoff E, Rutter M. The child and adolescent psychiatric assessment (CAPA). *Psychological medicine* 1995; **25**(04): 739-53.
5. American Psychiatric Association. Diagnostic and statistical manual of mental disorders, text revision (DSM-IV-TR): American Psychiatric Association; 2000.
6. Simonoff E, Pickles A, Charman T, Chandler S, Loucas T, Baird G. Psychiatric disorders in children with autism spectrum disorders: prevalence, comorbidity, and associated factors in a population-derived sample. *Journal of the American Academy of Child & Adolescent Psychiatry* 2008; **47**(8): 921-9.
7. Niarchou M, Zammit S, van Goozen SHM, et al. Psychopathology and cognition in children with 22q11.2 deletion syndrome. *British Journal of Psychiatry* 2014; **204**(1): 46-54.
8. Baker KD, Skuse DH. Adolescents and young adults with 22q11 deletion syndrome: psychopathology in an at-risk group. *British Journal of Psychiatry* 2005; **186**: 115-20.
9. Rutter M, Bailey A, Lord C. The social communication questionnaire: Manual: Western Psychological Services; 2003.
10. Berument SK, Rutter M, Lord C, Pickles A, Bailey A. Autism screening questionnaire: diagnostic validity. *British Journal of Psychiatry* 1999; **175**.
11. Wilson BN, Crawford SG, Green D, Roberts G, Aylott A, Kaplan BJ. Psychometric properties of the revised Developmental Coordination Disorder Questionnaire. *Physical & occupational therapy in pediatrics* 2009; **29**(2): 182-202.
12. Goodman R. The Strengths and Difficulties Questionnaire: a research note. *Journal of child psychology and psychiatry, and allied disciplines* 1997; **38**(5): 581-6.
13. Shaffer D, Gould MS, Brasic J, et al. A children's global assessment scale (CGAS). *Archives of General psychiatry* 1983; **40**(11): 1228-31.

### Supplementary Table 1



| Domain | CNV | 1q21.1 distal deletion | 1q21.1 distal duplication | 1q21.1 proximal duplication | 2p16.3 deletion | 9q34 deletion | 15q11.2 deletion | 15q13.3 deletion | 15q13.3 duplication | 16p11.2 distal deletion | 16p11.2 proximal deletion | 16p11.2 proximal duplication | 22q11.2 deletion | 22q11.2 duplication | Combined deletion group | Combined duplication group | Control siblings |
|---|---|---|---|---|---|---|---|---|---|---|---|---|---|---|---|---|---|
| FSIQ | N | 20 | 21 | 11 | 12 | 9 | 28 | 19 | 11 | 11 | 42 | 18 | 16 | 19 | 157 | 80 | 100 |
| | orginal score | 81.05 | 87.62 | 82.00 | 79.58 | 59.11 | 81.79 | 74.47 | 88.18 | 80.91 | 77.36 | 77.28 | 76.06 | 87.63 | 77.51 | 84.60 | 96.97 |
| | orginal score SD | 13.32 | 17.96 | 12.04 | 16.56 | 11.61 | 10.92 | 12.90 | 18.95 | 10.93 | 10.49 | 12.40 | 8.24 | 17.46 | 12.56 | 16.36 | 13.05 |
| | zscore | -1.37 | -0.90 | -1.30 | -1.58 | -3.35 | -1.32 | -1.94 | -0.78 | -1.32 | -1.68 | -1.65 | -1.76 | -0.88 | -1.68 | -1.10 | 0.00 |
| | zscore SD | 1.14 | 1.45 | 1.02 | 1.32 | 0.97 | 0.92 | 1.12 | 1.56 | 0.87 | 0.90 | 1.07 | 0.83 | 1.42 | 1.08 | 1.34 | 1.00 |
| | p-value (contrasted to controls) | 1.76E-05 | 1.96E-02 | 2.28E-02 | 1.30E-04 | 1.11E-16 | 8.78E-08 | 3.59E-12 | 4.61E-01 | 8.55E-03 | 5.55E-16 | 2.15E-07 | 1.91E-09 | 3.19E-02 | <1.00E-15 | 1.56E-11 | - |
| PIQ | N | 20 | 21 | 11 | 11 | 9 | 29 | 19 | 11 | 11 | 43 | 18 | 16 | 19 | 158 | 80 | 99 |
| | orginal score | 86.00 | 92.10 | 87.82 | 84.91 | 63.89 | 89.03 | 78.68 | 94.73 | 81.18 | 84.37 | 80.72 | 77.44 | 92.53 | 82.70 | 89.41 | 98.64 |
| | orginal score SD | 12.20 | 17.67 | 12.90 | 17.05 | 11.96 | 13.17 | 11.77 | 18.26 | 9.71 | 11.17 | 11.55 | 7.87 | 20.88 | 13.08 | 17.21 | 14.59 |
| | zscore | -0.96 | -0.59 | -0.85 | -1.13 | -2.58 | -0.77 | -1.48 | -0.32 | -1.25 | -1.07 | -1.29 | -1.56 | -0.57 | -1.20 | -0.74 | 0.00 |
| | zscore SD | 0.91 | 1.23 | 0.91 | 1.19 | 0.77 | 0.93 | 0.84 | 1.32 | 0.70 | 0.79 | 0.80 | 0.66 | 1.46 | 0.94 | 1.20 | 1.00 |
| | p-value (contrasted to controls) | 4.96E-03 | 3.15E-01 | 5.33E-01 | 2.09E-02 | 3.80E-13 | 7.26E-03 | 1.06E-07 | 9.85E-01 | 7.00E-03 | 4.22E-07 | 4.31E-05 | 4.65E-10 | 4.69E-01 | <1.00E-15 | 1.03E-06 | - |
| VIQ | N | 20 | 21 | 11 | 13 | 9 | 29 | 19 | 11 | 11 | 42 | 18 | 16 | 19 | 159 | 80 | 100 |
| | orginal score | 79.60 | 85.24 | 79.73 | 76.85 | 60.56 | 77.97 | 75.00 | 83.64 | 84.00 | 74.31 | 77.94 | 78.31 | 85.16 | 76.23 | 82.60 | 96.02 |
| | orginal score SD | 14.61 | 17.23 | 12.12 | 15.56 | 10.49 | 10.97 | 14.07 | 17.39 | 11.91 | 12.55 | 16.41 | 12.73 | 14.08 | 13.44 | 15.65 | 14.03 |
| | zscore | -1.32 | -0.92 | -1.30 | -1.60 | -2.97 | -1.44 | -1.69 | -0.99 | -0.90 | -1.76 | -1.45 | -1.41 | -0.87 | -1.60 | -1.09 | 0.00 |
| | zscore SD | 1.18 | 1.33 | 1.00 | 1.25 | 0.90 | 0.91 | 1.16 | 1.36 | 0.90 | 1.03 | 1.38 | 1.14 | 1.11 | 1.11 | 1.25 | 1.00 |
| | p-value (contrasted to controls) | 9.72E-05 | 1.51E-02 | 1.62E-02 | 1.68E-05 | 1.16E-13 | 1.91E-09 | 3.22E-08 | 1.76E-01 | 2.58E-01 | <1.00E-15 | 9.35E-06 | 2.76E-05 | 2.71E-02 | <1.00E-15 | 1.53E-11 | - |
| Non-verbal reasoning | N | 20 | 21 | 12 | 10 | 9 | 29 | 19 | 11 | 11 | 43 | 18 | 16 | 19 | 157 | 81 | 100 |
| | orginal score | 38.45 | 46.86 | 41.00 | 39.30 | 25.67 | 44.10 | 33.95 | 44.64 | 37.91 | 39.33 | 35.44 | 34.00 | 44.21 | 38.02 | 42.53 | 48.56 |
| | orginal score SD | 8.91 | 12.50 | 9.40 | 10.27 | 8.99 | 10.79 | 8.75 | 11.76 | 9.33 | 10.13 | 10.57 | 7.08 | 15.38 | 10.37 | 12.78 | 9.87 |
| | zscore | -1.15 | -0.34 | -0.93 | -1.16 | -2.26 | -0.56 | -1.54 | -0.42 | -1.12 | -1.04 | -1.35 | -1.61 | -0.55 | -1.17 | -0.71 | 0.00 |
| | zscore SD | 0.98 | 1.29 | 0.98 | 1.11 | 0.66 | 1.07 | 0.72 | 1.22 | 0.98 | 1.00 | 1.02 | 0.72 | 1.56 | 1.02 | 1.29 | 1.00 |
| | p-value (contrasted to controls) | 8.98E-04 | 9.76E-01 | 3.91E-01 | 2.57E-02 | 2.79E-08 | 2.68E-01 | 2.44E-07 | 9.69E-01 | 7.12E-02 | 3.46E-06 | 4.50E-05 | 3.85E-08 | 5.69E-01 | <1.00E-15 | 1.02E-05 | - |
| Perceptual organisation | N | 20 | 21 | 12 | 9 | 9 | 29 | 19 | 11 | 11 | 43 | 18 | 16 | 19 | 156 | 81 | 99 |
| | orginal score | 43.40 | 43.14 | 44.42 | 41.11 | 27.44 | 41.76 | 37.32 | 47.18 | 37.18 | 40.16 | 40.22 | 36.69 | 42.89 | 39.28 | 43.17 | 49.15 |
| | orginal score SD | 10.99 | 11.11 | 9.31 | 12.59 | 7.60 | 9.44 | 8.59 | 13.75 | 5.98 | 7.02 | 11.46 | 6.65 | 12.72 | 9.16 | 11.63 | 11.44 |
| | zscore | -0.56 | -0.60 | -0.46 | -0.85 | -2.32 | -0.73 | -1.17 | -0.25 | -1.09 | -0.82 | -0.87 | -1.15 | -0.68 | -0.95 | -0.61 | 0.00 |
| | zscore SD | 0.99 | 1.04 | 0.82 | 1.10 | 0.87 | 0.90 | 0.93 | 1.25 | 0.56 | 0.71 | 1.05 | 0.74 | 1.15 | 0.91 | 1.06 | 1.00 |
| | p-value (contrasted to controls) | 3.74E-01 | 2.33E-01 | 9.77E-01 | 3.04E-01 | 5.06E-10 | 1.02E-02 | 8.56E-05 | 9.98E-01 | 1.95E-02 | 1.13E-04 | 3.68E-02 | 3.37E-05 | 1.88E-01 | <1.00E-15 | 2.87E-05 | - |
| Verbal knowledge | N | 20 | 21 | 11 | 11 | 9 | 29 | 19 | 11 | 11 | 42 | 18 | 16 | 19 | 157 | 80 | 100 |
| | orginal score | 32.50 | 35.95 | 30.64 | 28.64 | 22.22 | 31.62 | 30.16 | 36.09 | 35.55 | 29.81 | 31.44 | 33.31 | 37.63 | 30.77 | 34.63 | 44.73 |
| | orginal score SD | 10.30 | 12.41 | 9.24 | 9.79 | 5.07 | 9.33 | 10.65 | 12.67 | 6.92 | 8.59 | 10.71 | 10.08 | 10.08 | 9.43 | 11.20 | 11.19 |
| | zscore | -1.20 | -0.88 | -1.39 | -1.62 | -2.26 | -1.29 | -1.45 | -0.85 | -0.83 | -1.46 | -1.30 | -1.10 | -0.68 | -1.37 | -0.99 | 0.00 |
| | zscore SD | 1.07 | 1.22 | 0.98 | 0.98 | 0.53 | 0.96 | 1.08 | 1.21 | 0.65 | 0.89 | 1.09 | 1.05 | 1.01 | 0.97 | 1.11 | 1.00 |
| | p-value (contrasted to controls) | 1.52E-04 | 6.22E-03 | 2.00E-03 | 3.06E-05 | 8.34E-10 | 9.82E-09 | 6.96E-08 | 3.60E-01 | 2.97E-01 | 2.44E-15 | 2.63E-05 | 5.89E-04 | 1.42E-01 | <1.00E-15 | 2.95E-11 | - |
| Verbal reasoning | N | 20 | 21 | 12 | 11 | 9 | 29 | 19 | 11 | 11 | 42 | 18 | 16 | 19 | 157 | 81 | 100 |
| | orginal score | 39.00 | 43.43 | 40.00 | 36.91 | 25.11 | 38.07 | 35.32 | 42.36 | 42.55 | 33.79 | 38.22 | 36.94 | 40.53 | 36.08 | 40.94 | 49.61 |
| | orginal score SD | 12.01 | 12.50 | 10.69 | 14.22 | 10.39 | 10.36 | 11.58 | 13.09 | 11.28 | 11.06 | 12.89 | 9.93 | 13.27 | 11.57 | 12.44 | 10.50 |
| | zscore | -1.09 | -0.68 | -1.00 | -1.36 | -2.45 | -1.19 | -1.44 | -0.73 | -0.70 | -1.61 | -1.14 | -1.28 | -0.95 | -1.38 | -0.90 | 0.00 |
| | zscore SD | 1.22 | 1.22 | 1.07 | 1.41 | 0.99 | 1.03 | 1.14 | 1.27 | 1.07 | 1.10 | 1.30 | 1.05 | 1.31 | 1.15 | 1.23 | 1.00 |
| | p-value (contrasted to controls) | 3.01E-03 | 3.10E-01 | 1.06E-01 | 4.65E-03 | 1.55E-08 | 3.50E-05 | 1.06E-05 | 6.65E-01 | 7.35E-01 | 2.44E-14 | 3.72E-03 | 1.57E-03 | 2.25E-02 | <1.00E-15 | 1.47E-07 | - |
| Set-shifting | N | 17 | 17 | 11 | 9 | 6 | 24 | 14 | 11 | 10 | 35 | 15 | 14 | 15 | 129 | 69 | 87 |
| | orginal score | 103.65 | 106.71 | 92.55 | 100.67 | 76.00 | 91.79 | 87.50 | 109.91 | 94.60 | 94.74 | 93.47 | 88.43 | 95.40 | 93.43 | 99.62 | 112.20 |
| | orginal score SD | 25.23 | 29.55 | 30.23 | 34.75 | 25.88 | 22.09 | 24.12 | 27.84 | 23.86 | 26.43 | 28.93 | 14.99 | 20.74 | 24.81 | 27.65 | 24.37 |
| | zscore | -0.36 | -0.18 | -0.87 | -0.57 | -1.81 | -0.88 | -1.15 | -0.11 | -0.77 | -0.75 | -0.87 | -0.93 | -0.68 | -0.82 | -0.54 | 0.00 |
| | zscore SD | 1.04 | 1.30 | 1.29 | 1.48 | 1.28 | 1.01 | 1.19 | 1.15 | 1.01 | 1.17 | 1.17 | 0.73 | 0.89 | 1.12 | 1.18 | 1.00 |
| | p-value (contrasted to controls) | 9.90E-01 | 1.00E+00 | 4.09E-01 | 9.68E-01 | 2.18E-03 | 9.43E-03 | 6.51E-03 | 1.00E+00 | 3.05E-01 | 1.44E-02 | 4.88E-02 | 3.32E-02 | 2.48E-01 | 6.47E-10 | 5.52E-04 | - |
| Spatial working memory | N | 15 | 20 | 14 | 11 | 7 | 30 | 14 | 9 | 10 | 38 | 16 | 13 | 15 | 138 | 74 | 64 |
| | orginal score | -1.17 | -0.63 | -1.27 | -1.35 | -1.53 | -0.77 | -1.99 | -0.35 | -0.76 | -1.18 | -1.30 | -1.15 | -1.54 | -1.17 | -1.05 | -0.31 |
| | orginal score SD | 1.22 | 1.11 | 1.44 | 1.31 | 1.85 | 1.40 | 1.51 | 1.28 | 0.94 | 1.24 | 1.18 | 0.93 | 1.31 | 1.32 | 1.29 | 1.12 |
| | zscore | -0.89 | -0.46 | -0.96 | -1.07 | -0.99 | -0.49 | -1.43 | -0.04 | -0.49 | -0.88 | -0.94 | -0.86 | -1.17 | -0.84 | -0.75 | 0.00 |
| | zscore SD | 1.03 | 1.00 | 1.20 | 0.98 | 1.47 | 1.36 | 1.10 | 1.31 | 0.82 | 0.93 | 0.92 | 0.72 | 1.05 | 1.09 | 1.11 | 1.00 |
| | p-value (contrasted to controls) | 1.25E-01 | 8.93E-01 | 8.84E-02 | 7.19E-02 | 5.40E-01 | 5.80E-01 | 2.43E-04 | 1.00E+00 | 9.80E-01 | 2.24E-03 | 6.74E-02 | 2.83E-01 | 5.69E-03 | 2.83E-07 | 8.26E-05 | - |
| Spatial planning | N | 11 | 18 | 11 | 7 | 5 | 27 | 10 | 8 | 8 | 32 | 10 | 11 | 12 | 111 | 59 | 60 |
| | orginal score | -0.80 | -0.87 | -0.80 | -1.40 | -2.85 | -1.09 | -1.39 | -1.06 | -0.92 | -0.74 | -1.66 | -1.48 | -1.42 | -1.11 | -1.13 | -0.55 |
| | orginal score SD | 0.82 | 0.76 | 0.92 | 0.91 | 1.75 | 0.92 | 1.22 | 1.54 | 0.59 | 1.02 | 1.10 | 0.70 | 1.12 | 1.06 | 1.07 | 0.97 |
| | zscore | -0.30 | -0.45 | -0.31 | -0.89 | -1.83 | -0.58 | -0.76 | -0.41 | -0.42 | -0.23 | -1.02 | -0.98 | -0.85 | -0.57 | -0.59 | 0.00 |
| | zscore SD | 0.81 | 0.79 | 0.92 | 0.82 | 1.18 | 0.78 | 0.99 | 1.37 | 0.57 | 1.05 | 0.86 | 0.63 | 0.93 | 0.94 | 0.95 | 1.00 |
| | p-value (contrasted to controls) | 9.99E-01 | 9.00E-01 | 9.99E-01 | 4.75E-01 | 1.89E-03 | 2.85E-01 | 4.59E-01 | 9.96E-01 | 9.94E-01 | 9.98E-01 | 7.19E-02 | 7.11E-02 | 1.57E-01 | 4.19E-04 | 1.68E-03 | - |
| Sustained attention | N | 14 | 12 | 11 | 8 | 3 | 21 | 9 | 6 | 9 | 29 | 11 | 11 | 11 | 104 | 51 | 57 |
| | orginal score | -2.14 | -1.49 | -2.29 | -4.54 | -2.42 | -1.99 | -1.95 | -1.36 | -2.63 | -1.87 | -1.90 | -2.25 | -0.91 | -2.26 | -1.61 | -0.22 |
| | orginal score SD | 2.74 | 2.91 | 3.12 | 4.89 | 3.46 | 2.77 | 2.62 | 1.62 | 3.41 | 1.73 | 2.19 | 3.00 | 1.91 | 2.81 | 2.45 | 1.15 |
| | zscore | -0.87 | -0.37 | -0.77 | -1.32 | -2.42 | -0.78 | -1.34 | -1.32 | -1.33 | -1.16 | -1.36 | -1.14 | -0.29 | -1.12 | -0.77 | 0.00 |
| | zscore SD | 1.23 | 1.36 | 1.34 | 1.12 | 2.84 | 1.44 | 1.42 | 0.49 | 1.10 | 1.04 | 0.91 | 1.39 | 1.22 | 1.29 | 1.20 | 1.00 |
| | p-value (contrasted to controls) | 4.31E-01 | 9.99E-01 | 7.89E-01 | 1.59E-01 | 5.07E-02 | 3.22E-01 | 8.69E-02 | 3.49E-01 | 9.10E-02 | 1.86E-03 | 3.66E-02 | 1.57E-01 | 1.00E+00 | 8.10E-03 | 3.91E-01 | - |
| Processing speed | N | 14 | 19 | 13 | 10 | 4 | 27 | 13 | 8 | 9 | 34 | 16 | 11 | 16 | 122 | 72 | 62 |
| | orginal score | -1.11 | -0.36 | -0.41 | -1.87 | -7.41 | -1.02 | -2.88 | -0.48 | -0.79 | -0.52 | -1.13 | 0.58 | -0.42 | -1.21 | -0.56 | -0.14 |
| | orginal score SD | 2.63 | 1.20 | 1.83 | 1.94 | 12.55 | 1.94 | 3.65 | 1.36 | 2.17 | 1.27 | 2.52 | 0.78 | 0.96 | 3.15 | 1.66 | 1.20 |
| | zscore | -0.45 | -0.11 | -0.04 | -1.08 | -1.48 | -0.51 | -1.35 | -0.29 | -0.27 | -0.28 | -0.47 | 0.65 | -0.26 | -0.48 | -0.23 | 0.00 |
| | zscore SD | 1.05 | 0.99 | 1.16 | 0.76 | 1.19 | 1.12 | 1.03 | 1.01 | 1.42 | 0.93 | 1.18 | 0.70 | 0.78 | 1.12 | 1.01 | 1.00 |
| | p-value (contrasted to controls) | 9.71E-01 | 1.00E+00 | 1.00E+00 | 1.01E-01 | 3.45E-01 | 5.43E-01 | 9.99E-04 | 1.00E+00 | 1.00E+00 | 9.95E-01 | 9.07E-01 | 9.45E-01 | 1.00E+00 | 3.91E-01 | 8.10E-03 | - |
| Total CAPA symptom count | N | 16 | 19 | 12 | 13 | 10 | 30 | 20 | 10 | 12 | 44 | 14 | 16 | 17 | 161 | 72 | 75 |
| | orginal score | 26.00 | 21.21 | 18.67 | 23.85 | 12.00 | 23.50 | 19.15 | 25.10 | 26.00 | 15.68 | 34.07 | 8.56 | 20.24 | 19.09 | 23.60 | 7.92 |
| | orginal score SD | 14.17 | 10.05 | 8.28 | 15.80 | 7.23 | 11.59 | 11.73 | 15.57 | 13.74 | 10.49 | 21.44 | 7.73 | 14.59 | 12.67 | 15.14 | 9.27 |
| | zscore | -1.71 | -1.49 | -1.22 | -1.41 | -0.65 | -1.50 | -1.12 | -1.59 | -1.77 | -0.84 | -2.16 | -0.10 | -1.12 | -1.11 | -1.50 | 0.00 |
| | zscore SD | 1.11 | 0.95 | 0.77 | 1.29 | 0.91 | 1.01 | 1.12 | 1.29 | 1.31 | 1.10 | 1.24 | 0.89 | 1.29 | 1.17 | 1.15 | 1.00 |
| | p-value (contrasted to controls) | 1.27E-07 | 1.13E-05 | 1.93E-02 | 3.46E-04 | 8.76E-01 | 5.85E-10 | 2.13E-03 | 4.84E-04 | 1.11E-05 | 2.78E-03 | 2.41E-10 | 1.00E+00 | 1.57E-03 | 1.75E-12 | 4.66E-15 | - |
| Anxiety CAPA subscale | N | 16 | 19 | 12 | 13 | 10 | 30 | 20 | 10 | 12 | 44 | 14 | 16 | 17 | 161 | 72 | 75 |
| | orginal score | 4.63 | 4.68 | 5.42 | 6.31 | 1.80 | 5.30 | 2.40 | 6.20 | 4.67 | 2.84 | 8.71 | 0.31 | 4.06 | 3.52 | 5.65 | 1.36 |
| | orginal score SD | 5.26 | 4.16 | 5.43 | 8.43 | 2.74 | 4.65 | 3.42 | 6.89 | 4.77 | 3.58 | 7.69 | 0.87 | 5.26 | 4.66 | 5.90 | 2.85 |
| | zscore | -0.98 | -1.20 | -1.30 | -1.10 | -0.26 | -1.27 | -0.35 | -1.30 | -1.10 | -0.53 | -1.92 | 0.39 | -0.76 | -0.67 | -1.26 | 0.00 |
| | zscore SD | 1.55 | 1.36 | 1.44 | 1.73 | 1.14 | 1.40 | 1.21 | 1.71 | 1.48 | 1.27 | 1.60 | 0.40 | 1.57 | 1.38 | 1.53 | 1.00 |
| | p-value (contrasted to controls) | 1.34E-01 | 2.74E-02 | 6.21E-02 | 1.23E-01 | 1.00E+00 | 1.64E-04 | 9.96E-01 | 1.00E-01 | 2.37E-01 | 5.62E-01 | 5.14E-05 | 9.99E-01 | 4.15E-01 | 6.37E-04 | 2.10E-08 | - |
| ADHD CAPA subscale | N | 16 | 19 | 12 | 13 | 10 | 30 | 20 | 10 | 12 | 44 | 14 | 16 | 17 | 161 | 72 | 75 |
| | orginal score | 8.94 | 9.21 | 4.92 | 9.08 | 5.50 | 8.73 | 8.00 | 9.10 | 9.83 | 5.98 | 10.86 | 4.25 | 7.88 | 7.37 | 8.49 | 2.44 |
| | orginal score SD | 4.23 | 4.01 | 3.06 | 4.15 | 3.54 | 4.36 | 4.99 | 5.72 | 5.75 | 4.63 | 4.88 | 3.89 | 6.16 | 4.76 | 5.11 | 4.06 |
| | zscore | -1.46 | -1.64 | -0.51 | -1.43 | -0.89 | -1.36 | -1.24 | -1.50 | -1.79 | -0.78 | -2.09 | -0.40 | -1.16 | -1.11 | -1.41 | 0.00 |
| | zscore SD | 1.10 | 1.01 | 0.70 | 1.15 | 0.91 | 1.13 | 1.26 | 1.35 | 1.57 | 1.16 | 1.22 | 0.97 | 1.54 | 1.21 | 1.28 | 1.00 |
| | p-value (contrasted to controls) | 1.83E-04 | 7.68E-07 | 9.79E-01 | 1.16E-03 | 6.16E-01 | 7.80E-07 | 7.02E-04 | 3.55E-03 | 2.34E-05 | 2.23E-02 | 3.37E-08 | 9.90E-01 | 4.30E-03 | 3.80E-11 | 1.13E-12 | - |
| Mood CAPA subscale | N | 16 | 19 | 12 | 13 | 10 | 30 | 20 | 10 | 12 | 44 | 14 | 16 | 17 | 161 | 72 | 75 |
| | orginal score | 3.81 | 1.84 | 2.58 | 2.31 | 1.20 | 2.50 | 2.20 | 2.40 | 4.00 | 2.36 | 6.14 | 1.00 | 2.24 | 2.42 | 2.97 | 1.21 |
| | orginal score SD | 3.75 | 1.80 | 3.20 | 3.28 | 2.15 | 2.35 | 2.50 | 2.17 | 4.16 | 3.71 | 6.55 | 1.26 | 2.66 | 3.14 | 3.87 | 1.80 |
| | zscore | -1.13 | -0.50 | -0.69 | -0.39 | 0.07 | -0.72 | -0.49 | -0.72 | -1.10 | -0.44 | -1.36 | 0.03 | -0.52 | -0.53 | -0.73 | 0.00 |
| | zscore SD | 1.19 | 0.97 | 1.00 | 1.21 | 1.06 | 1.14 | 1.17 | 1.23 | 1.24 | 1.09 | 1.18 | 0.92 | 1.15 | 1.15 | 1.12 | 1.00 |
| | p-value (contrasted to controls) | 6.47E-03 | 9.23E-01 | 6.84E-01 | 9.92E-01 | 1.00E+00 | 7.04E-02 | 8.10E-01 | 7.43E-01 | 5.10E-02 | 5.99E-01 | 1.43E-03 | 1.00E+00 | 7.78E-01 | 1.32E-03 | 1.94E-04 | - |
| OCD CAPA subscale | N | 16 | 19 | 12 | 13 | 10 | 30 | 20 | 10 | 12 | 44 | 14 | 16 | 17 | 161 | 72 | 75 |
| | orginal score | 0.13 | 0.26 | 0.17 | 0.38 | 0.40 | 0.17 | 0.20 | 0.30 | 0.58 | 0.18 | 0.43 | 0.00 | 0.35 | 0.22 | 0.31 | 0.05 |
| | orginal score SD | 0.34 | 0.56 | 0.39 | 0.96 | 0.52 | 0.38 | 0.41 | 0.48 | 0.90 | 0.45 | 0.65 | 0.00 | 1.22 | 0.52 | 0.74 | 0.23 |
| | zscore | -0.32 | -0.90 | -0.51 | -0.80 | -1.50 | -0.42 | -0.55 | -1.05 | -1.79 | -0.50 | -1.41 | 0.16 | -0.49 | -0.59 | -0.86 | 0.00 |
| | zscore SD | 1.51 | 2.06 | 1.56 | 2.55 | 2.24 | 1.58 | 1.66 | 2.01 | 2.66 | 1.70 | 2.27 | 0.20 | 2.16 | 1.81 | 2.02 | 1.00 |
| | p-value (contrasted to controls) | 1.00E+00 | 6.77E-01 | 9.98E-01 | 9.61E-01 | 2.88E-01 | 9.96E-01 | 9.94E-01 | 8.27E-01 | 4.39E-02 | 9.63E-01 | 2.78E-01 | 1.00E+00 | 9.90E-01 | 2.34E-02 | 2.73E-03 | - |
| ODD CAPA subscale | N | 16 | 19 | 12 | 13 | 10 | 30 | 20 | 10 | 12 | 44 | 14 | 16 | 17 | 161 | 72 | 75 |
| | orginal score | 5.88 | 3.37 | 3.25 | 3.92 | 1.20 | 4.23 | 4.55 | 5.20 | 5.17 | 2.91 | 5.43 | 1.81 | 4.18 | 3.69 | 4.19 | 1.97 |
| | orginal score SD | 3.12 | 2.54 | 1.82 | 3.95 | 1.03 | 3.01 | 3.36 | 3.58 | 4.11 | 2.77 | 3.50 | 2.14 | 4.26 | 3.24 | 3.30 | 2.22 |
| | zscore | -1.49 | -0.62 | -0.64 | -0.67 | 0.28 | -0.88 | -0.98 | -1.20 | -1.13 | -0.37 | -1.33 | 0.09 | -0.71 | -0.65 | -0.86 | 0.00 |
| | zscore SD | 0.98 | 1.07 | 0.81 | 1.12 | 0.64 | 1.14 | 1.20 | 1.26 | 1.44 | 1.10 | 1.12 | 0.97 | 1.28 | 1.19 | 1.13 | 1.00 |
| | p-value (contrasted to controls) | 3.70E-05 | 6.57E-01 | 9.11E-01 | 5.62E-01 | 1.00E+00 | 5.80E-03 | 2.11E-02 | 3.28E-02 | 3.42E-02 | 8.45E-01 | 3.98E-03 | 1.00E+00 | 2.85E-01 | 2.36E-04 | 4.05E-05 | - |
| Sleep CAPA subscale | N | 16 | 19 | 12 | 13 | 10 | 30 | 20 | 10 | 12 | 44 | 14 | 16 | 17 | 161 | 72 | 75 |
| | orginal score | 2.63 | 1.84 | 2.33 | 1.85 | 1.90 | 2.57 | 1.80 | 1.90 | 1.75 | 1.41 | 2.50 | 1.19 | 1.53 | 1.86 | 1.99 | 0.88 |
| | orginal score SD | 2.78 | 1.89 | 1.97 | 2.03 | 1.20 | 2.19 | 1.74 | 1.85 | 1.29 | 1.63 | 2.41 | 1.47 | 1.12 | 1.91 | 1.85 | 1.16 |
| | zscore | -1.02 | -0.77 | -1.06 | -0.53 | -0.99 | -1.05 | -0.58 | -0.69 | -0.70 | -0.35 | -1.07 | -0.26 | -0.47 | -0.64 | -0.79 | 0.00 |
| | zscore SD | 1.72 | 1.38 | 1.44 | 1.56 | 1.18 | 1.44 | 1.27 | 1.26 | 1.03 | 1.25 | 1.54 | 1.15 | 0.99 | 1.35 | 1.31 | 1.00 |
| | p-value (contrasted to controls) | 1.35E-01 | 7.95E-01 | 2.28E-01 | 9.43E-01 | 5.46E-01 | 8.08E-03 | 8.57E-01 | 9.38E-01 | 7.64E-01 | 9.85E-01 | 2.47E-01 | 1.00E+00 | 9.86E-01 | 5.24E-04 | 9.62E-04 | - |
| Subclinical psychotic experiences (child CAPA) | N | 19 | 20 | 12 | 13 | 10 | 33 | 20 | 11 | 12 | 44 | 16 | 16 | 18 | 167 | 77 | 83 |
| | orginal score | 0.16 | 0.15 | 0.42 | 0.00 | 0.00 | 0.03 | 0.00 | 0.00 | 0.08 | 0.27 | 1.50 | 0.00 | 0.06 | 0.10 | 0.43 | 0.02 |
| | orginal score SD | 0.69 | 0.67 | 1.00 | 0.00 | 0.00 | 0.17 | 0.00 | 0.00 | 0.29 | 1.13 | 3.54 | 0.00 | 0.24 | 0.64 | 1.75 | 0.22 |
| | zscore | -0.55 | -0.63 | -1.60 | 0.03 | 0.15 | -0.13 | 0.10 | 0.06 | -0.36 | -0.77 | -3.10 | 0.00 | -0.30 | -0.29 | -1.12 | 0.00 |
| | zscore SD | 2.43 | 2.41 | 3.57 | 0.29 | 0.35 | 0.85 | 0.36 | 0.29 | 1.52 | 2.69 | 4.81 | 0.25 | 1.28 | 1.72 | 3.09 | 1.00 |
| | p-value (contrasted to controls) | 9.97E-01 | 9.89E-01 | 2.79E-01 | 1.00E+00 | 1.00E+00 | 1.00E+00 | 1.00E+00 | 1.00E+00 | 1.00E+00 | 6.65E-01 | 8.06E-07 | 1.00E+00 | 1.00E+00 | 4.98E-01 | 1.13E-03 | - |
| General functioning | N | 13 | 9 | 6 | 9 | 6 | 22 | 10 | 6 | 3 | 20 | 9 | 13 | 12 | 96 | 42 | 56 |
| | orginal score | 59.77 | 62.33 | 57.00 | 54.00 | 37.17 | 52.55 | 56.60 | 50.83 | 64.67 | 63.25 | 56.33 | 60.92 | 68.17 | 56.86 | 60.31 | 78.45 |
| | orginal score SD | 19.01 | 13.84 | 12.10 | 17.85 | 18.48 | 19.34 | 11.97 | 24.42 | 8.02 | 10.49 | 8.05 | 21.06 | 8.84 | 17.60 | 14.10 | 11.92 |
| | zscore | -1.20 | -1.09 | -1.56 | -1.52 | -2.65 | -1.60 | -1.47 | -1.70 | -0.95 | -1.10 | -1.62 | -1.13 | -0.76 | -1.40 | -1.26 | 0.00 |
| | zscore SD | 0.98 | 0.92 | 0.73 | 0.91 | 0.95 | 0.94 | 0.69 | 1.45 | 0.81 | 0.84 | 0.70 | 1.18 | 0.73 | 0.98 | 0.93 | 1.00 |
| | p-value (contrasted to controls) | 3.36E-04 | 1.83E-02 | 3.86E-03 | 7.77E-05 | 1.30E-09 | 7.75E-12 | 6.90E-05 | 4.26E-04 | 7.47E-01 | 6.99E-05 | 2.11E-05 | 1.94E-03 | 1.81E-01 | <1.00E-15 | 1.94E-11 | - |
| Social functioning | N | 11 | 9 | 6 | 9 | 6 | 22 | 8 | 4 | 3 | 18 | 8 | 13 | 11 | 90 | 38 | 56 |
| | orginal score | 58.27 | 62.00 | 53.83 | 48.67 | 33.83 | 51.45 | 53.13 | 44.00 | 55.33 | 61.22 | 54.13 | 58.85 | 68.36 | 54.13 | 59.00 | 79.16 |
| | orginal score SD | 18.37 | 14.30 | 12.86 | 19.89 | 22.61 | 20.62 | 20.32 | 19.71 | 5.03 | 10.01 | 12.88 | 17.63 | 11.47 | 18.57 | 15.07 | 10.87 |
| | zscore | -1.37 | -1.19 | -1.89 | -1.91 | -2.92 | -1.73 | -1.61 | -2.32 | -1.71 | -1.36 | -1.85 | -1.44 | -0.81 | -1.66 | -1.45 | 0.00 |
| | zscore SD | 0.92 | 1.00 | 0.88 | 0.95 | 0.94 | 0.99 | 0.95 | 0.90 | 0.24 | 0.89 | 1.02 | 0.94 | 1.05 | 0.98 | 1.08 | 1.00 |
| | p-value (contrasted to controls) | 8.24E-05 | 5.15E-03 | 8.31E-05 | 1.62E-08 | 9.06E-12 | 2.75E-14 | 5.76E-05 | 3.73E-05 | 3.73E-02 | 3.16E-07 | 3.50E-06 | 5.38E-06 | 1.55E-01 | <1.00E-15 | 7.53E-13 | - |
| ASD traits | N | 13 | 20 | 13 | 13 | 8 | 31 | 18 | 10 | 11 | 39 | 17 | 16 | 18 | 149 | 78 | 86 |
| | orginal score | 18.15 | 22.40 | 15.54 | 22.23 | 19.75 | 19.13 | 17.94 | 19.90 | 21.73 | 15.28 | 17.88 | 12.63 | 17.94 | 17.69 | 18.92 | 4.63 |
| | orginal score SD | 6.49 | 7.34 | 7.94 | 7.55 | 8.38 | 8.48 | 8.83 | 8.52 | 7.36 | 7.58 | 6.00 | 5.98 | 9.42 | 8.06 | 8.02 | 5.67 |
| | zscore | -2.23 | -3.00 | -1.81 | -2.83 | -2.71 | -2.32 | -2.13 | -2.45 | -2.83 | -1.74 | -2.19 | -1.36 | -2.16 | -2.14 | -2.36 | 0.00 |
| | zscore SD | 1.18 | 1.24 | 1.31 | 1.33 | 1.75 | 1.44 | 1.52 | 1.37 | 1.17 | 1.31 | 1.04 | 1.08 | 1.70 | 1.40 | 1.38 | 1.00 |
| | p-value (contrasted to controls) | 4.03E-08 | 0.00E+00 | 4.90E-05 | 2.44E-14 | 6.14E-07 | 0.00E+00 | 4.54E-11 | 2.21E-07 | 4.85E-12 | 4.24E-14 | 2.91E-10 | 6.33E-04 | 1.00E-11 | <1.00E-15 | <1.00E-15 | - |
| Motor coordination | N | 13 | 19 | 13 | 13 | 8 | 32 | 17 | 10 | 11 | 38 | 17 | 17 | 18 | 149 | 77 | 86 |
| | orginal score | 27.77 | 30.00 | 41.31 | 34.23 | 23.63 | 33.31 | 33.65 | 29.20 | 31.55 | 34.55 | 34.24 | 42.88 | 35.33 | 33.70 | 33.99 | 62.51 |
| | orginal score SD | 8.22 | 10.93 | 11.03 | 13.70 | 5.26 | 11.46 | 11.67 | 16.17 | 14.71 | 16.18 | 11.46 | 13.50 | 12.82 | 13.52 | 12.63 | 14.82 |
| | zscore | -2.44 | -2.29 | -1.17 | -1.73 | -3.14 | -1.79 | -1.77 | -2.41 | -2.17 | -1.89 | -1.82 | -1.18 | -1.69 | -1.90 | -1.87 | 0.00 |
| | zscore SD | 1.07 | 1.29 | 0.88 | 1.23 | 1.00 | 1.08 | 1.08 | 1.55 | 1.33 | 1.30 | 1.06 | 1.03 | 1.32 | 1.21 | 1.27 | 1.00 |
| | p-value (contrasted to controls) | 3.69E-12 | 2.78E-15 | 2.09E-02 | 3.35E-06 | 2.95E-12 | 6.66E-15 | 5.27E-08 | 5.99E-09 | 1.32E-08 | <1.00E-15 | 4.91E-08 | 1.99E-03 | 1.10E-08 | <1.00E-15 | <1.00E-15 | - |
| SDQ total (caregiver report) | N | 13 | 20 | 13 | 13 | 8 | 32 | 18 | 10 | 11 | 39 | 17 | 17 | 18 | 151 | 78 | 88 |
| | orginal score | 23.92 | 22.40 | 18.62 | 22.38 | 15.88 | 22.47 | 19.72 | 22.30 | 24.73 | 16.92 | 23.76 | 15.53 | 21.22 | 19.86 | 21.78 | 8.93 |
| | orginal score SD | 6.49 | 5.73 | 6.33 | 4.23 | 5.03 | 7.01 | 6.37 | 8.29 | 5.48 | 6.20 | 6.46 | 5.14 | 7.30 | 6.76 | 6.75 | 7.08 |
| | zscore | -2.12 | -1.87 | -1.30 | -1.79 | -1.02 | -1.85 | -1.44 | -1.84 | -2.23 | -1.05 | -2.11 | -0.88 | -1.69 | -1.49 | -1.78 | 0.00 |
| | zscore SD | 1.10 | 0.84 | 0.89 | 0.70 | 0.70 | 1.11 | 0.95 | 1.30 | 0.92 | 0.94 | 1.09 | 0.75 | 1.16 | 1.04 | 1.05 | 1.00 |
| | p-value (contrasted to controls) | 7.85E-13 | 3.64E-14 | 2.50E-04 | 1.37E-09 | 3.39E-01 | <1.00E-15 | 2.98E-08 | 4.95E-07 | 2.51E-12 | 6.80E-08 | 3.33E-15 | 2.29E-02 | 6.61E-11 | <1.00E-15 | <1.00E-15 | - |
| Conduct SDQ subscale (caregiver report) | N | 13 | 20 | 13 | 13 | 8 | 32 | 18 | 10 | 11 | 39 | 17 | 17 | 18 | 151 | 78 | 88 |
| | orginal score | 5.23 | 3.95 | 3.69 | 3.23 | 1.13 | 4.75 | 3.44 | 4.00 | 4.64 | 2.74 | 4.76 | 1.82 | 4.22 | 3.46 | 4.15 | 2.24 |
| | orginal score SD | 2.45 | 2.11 | 1.75 | 2.65 | 0.83 | 2.78 | 2.53 | 2.94 | 2.77 | 2.12 | 2.86 | 2.07 | 2.82 | 2.64 | 2.49 | 2.25 |
| | zscore | -1.24 | -0.74 | -0.67 | -0.38 | 0.44 | -1.01 | -0.51 | -0.69 | -1.02 | -0.22 | -1.06 | 0.23 | -0.81 | -0.50 | -0.81 | 0.00 |
| | zscore SD | 0.90 | 0.94 | 0.84 | 1.09 | 0.55 | 1.07 | 1.04 | 1.33 | 1.14 | 0.95 | 1.04 | 0.97 | 1.17 | 1.10 | 1.04 | 1.00 |
| | p-value (contrasted to controls) | 2.51E-03 | 7.89E-02 | 7.14E-01 | 9.72E-01 | 9.92E-01 | 1.33E-05 | 6.30E-01 | 5.92E-01 | 4.88E-02 | 9.92E-01 | 4.82E-03 | 9.99E-01 | 4.18E-02 | 1.15E-03 | 4.13E-06 | - |
| Emotional SDQ subscale (caregiver report) | N | 13 | 20 | 13 | 13 | 8 | 32 | 18 | 10 | 11 | 39 | 17 | 17 | 18 | 151 | 78 | 88 |
| | orginal score | 6.00 | 4.95 | 5.23 | 5.69 | 3.13 | 4.72 | 4.83 | 5.50 | 5.36 | 4.05 | 6.12 | 3.24 | 5.33 | 4.55 | 5.41 | 2.84 |
| | orginal score SD | 2.45 | 2.67 | 2.62 | 2.78 | 2.36 | 2.98 | 2.64 | 2.72 | 2.54 | 2.67 | 2.85 | 2.39 | 2.99 | 2.75 | 2.74 | 2.66 |
| | zscore | -1.23 | -0.82 | -0.93 | -1.14 | -0.07 | -0.76 | -0.79 | -1.06 | -0.98 | -0.48 | -1.28 | -0.15 | -0.99 | -0.68 | -1.01 | 0.00 |
| | zscore SD | 0.95 | 1.03 | 1.01 | 1.05 | 0.95 | 1.14 | 1.00 | 1.04 | 0.95 | 1.01 | 1.08 | 0.88 | 1.14 | 1.05 | 1.05 | 1.00 |
| | p-value (contrasted to controls) | 2.01E-03 | 4.49E-02 | 7.92E-02 | 4.50E-03 | 1.00E+00 | 6.44E-03 | 7.43E-02 | 5.19E-02 | 8.95E-02 | 2.91E-01 | 9.84E-05 | 1.00E+00 | 6.10E-03 | 1.82E-06 | 4.01E-10 | - |
| Hyperactivity SDQ subscale (caregiver report) | N | 13 | 20 | 13 | 13 | 8 | 32 | 18 | 10 | 11 | 39 | 17 | 17 | 18 | 151 | 78 | 88 |
| | orginal score | 8.15 | 8.05 | 5.92 | 7.85 | 7.25 | 8.25 | 7.11 | 7.70 | 8.64 | 5.82 | 7.24 | 6.76 | 7.67 | 7.25 | 7.38 | 2.11 |
| | orginal score SD | 2.19 | 2.26 | 3.20 | 2.67 | 1.83 | 2.13 | 2.30 | 2.71 | 1.69 | 2.56 | 2.56 | 2.49 | 2.83 | 2.49 | 2.71 | 2.34 |
| | zscore | -2.73 | -2.70 | -1.55 | -2.44 | -2.64 | -2.68 | -2.07 | -2.40 | -3.09 | -1.46 | -2.32 | -2.10 | -2.48 | -2.24 | -2.34 | 0.00 |
| | zscore SD | 1.28 | 1.25 | 1.29 | 1.60 | 0.85 | 1.31 | 1.27 | 1.56 | 1.05 | 1.23 | 1.47 | 1.40 | 1.63 | 1.37 | 1.45 | 1.00 |
| | p-value (contrasted to controls) | 1.36E-13 | <1.00E-15 | 6.24E-04 | 2.68E-11 | 1.92E-06 | <1.00E-15 | 3.14E-10 | 4.70E-08 | 1.29E-14 | 1.41E-09 | 4.28E-11 | 1.29E-09 | 1.01E-14 | <1.00E-15 | <1.00E-15 | - |
| Peer SDQ subscale (caregiver report) | N | 13 | 20 | 13 | 13 | 8 | 32 | 18 | 10 | 11 | 39 | 17 | 17 | 18 | 151 | 78 | 88 |
| | orginal score | 4.54 | 5.45 | 3.77 | 5.62 | 4.38 | 4.75 | 4.33 | 5.10 | 6.09 | 4.31 | 5.65 | 3.71 | 4.00 | 4.60 | 4.83 | 1.74 |
| | orginal score SD | 2.88 | 2.70 | 2.28 | 2.26 | 2.97 | 2.27 | 2.30 | 2.28 | 1.70 | 2.51 | 2.42 | 2.05 | 2.45 | 2.40 | 2.52 | 2.16 |
| | zscore | -1.36 | -1.83 | -1.01 | -1.81 | -1.23 | -1.41 | -1.19 | -1.58 | -2.02 | -1.21 | -1.81 | -0.96 | -1.06 | -1.35 | -1.48 | 0.00 |
| | zscore SD | 1.35 | 1.25 | 1.08 | 1.06 | 1.31 | 1.06 | 1.09 | 1.11 | 0.79 | 1.14 | 1.14 | 0.96 | 1.20 | 1.11 | 1.20 | 1.00 |
| | p-value (contrasted to controls) | 8.84E-04 | 3.32E-10 | 5.50E-02 | 1.05E-07 | 1.80E-01 | 6.79E-10 | 5.02E-04 | 2.80E-04 | 5.07E-08 | 1.75E-07 | 3.15E-09 | 3.45E-02 | 3.60E-03 | <1.00E-15 | <1.00E-15 | - |
| Prosocial SDQ subscale (caregiver report) | N | 13 | 20 | 13 | 13 | 8 | 32 | 18 | 10 | 11 | 39 | 17 | 17 | 18 | 151 | 78 | 88 |
| | orginal score | 5.85 | 5.00 | 6.54 | 5.69 | 6.88 | 4.56 | 6.00 | 4.60 | 5.00 | 6.74 | 5.24 | 7.06 | 6.28 | 5.94 | 5.55 | 7.93 |
| | orginal score SD | 2.19 | 1.89 | 1.98 | 2.69 | 3.14 | 2.88 | 2.33 | 1.90 | 3.10 | 2.41 | 2.33 | 2.30 | 1.99 | 2.70 | 2.11 | 2.16 |
| | zscore | -0.88 | -1.29 | -0.60 | -0.84 | -0.55 | -1.34 | -0.76 | -1.36 | -1.21 | -0.47 | -1.15 | -0.39 | -0.66 | -0.81 | -1.01 | 0.00 |
| | zscore SD | 0.91 | 0.90 | 0.83 | 1.14 | 1.39 | 1.26 | 1.07 | 0.86 | 1.30 | 1.08 | 0.98 | 0.91 | 0.95 | 1.16 | 0.94 | 1.00 |
| | p-value (contrasted to controls) | 1.39E-01 | 2.79E-05 | 7.54E-01 | 1.39E-01 | 9.88E-01 | 8.38E-10 | 7.44E-02 | 1.95E-03 | 1.38E-02 | 4.90E-01 | 2.62E-03 | 9.81E-01 | 2.29E-01 | 1.20E-08 | 5.00E-09 | - |
| SDQ total (teacher report) | N | 7 | 11 | 5 | 11 | 5 | 19 | 11 | 6 | 6 | 26 | 10 | 10 | 11 | 95 | 43 | 52 |
| | orginal score | 13.43 | 10.45 | 13.40 | 10.73 | 10.00 | 12.00 | 10.82 | 14.00 | 9.67 | 9.35 | 13.90 | 9.20 | 14.09 | 10.55 | 13.02 | 5.19 |
| | orginal score SD | 6.50 | 6.92 | 5.13 | 6.05 | 6.75 | 6.32 | 5.08 | 3.52 | 5.16 | 3.90 | 6.15 | 3.79 | 6.49 | 5.24 | 5.99 | 5.02 |
| | zscore | -1.31 | -0.90 | -1.31 | -0.88 | -0.95 | -1.16 | -0.90 | -1.46 | -0.80 | -0.78 | -1.56 | -0.89 | -1.50 | -0.94 | -1.33 | 0.00 |
| | zscore SD | 1.24 | 1.21 | 0.91 | 1.15 | 1.22 | 1.07 | 0.83 | 0.79 | 0.97 | 0.77 | 0.94 | 0.76 | 1.15 | 0.94 | 1.04 | 1.00 |
| | p-value (contrasted to controls) | 2.39E-02 | 1.85E-01 | 1.04E-01 | 1.53E-01 | 7.32E-01 | 4.27E-04 | 1.12E-01 | 1.59E-02 | 7.46E-01 | 3.54E-02 | 1.77E-04 | 3.62E-01 | 1.14E-04 | 2.27E-08 | 2.54E-11 | - |
| Conduct SDQ subscale (teacher report) | N | 7 | 11 | 5 | 11 | 5 | 19 | 11 | 6 | 6 | 26 | 10 | 10 | 11 | 95 | 43 | 52 |
| | orginal score | 2.29 | 1.82 | 2.20 | 1.36 | 1.00 | 1.47 | 1.73 | 2.67 | 0.67 | 0.88 | 1.90 | 1.00 | 2.36 | 1.26 | 2.14 | 0.81 |
| | orginal score SD | 2.06 | 2.09 | 2.28 | 1.29 | 1.22 | 1.61 | 1.49 | 2.50 | 1.03 | 1.14 | 2.28 | 1.41 | 2.91 | 1.42 | 2.35 | 1.44 |
| | zscore | -0.89 | -0.64 | -0.80 | -0.38 | -0.25 | -0.43 | -0.59 | -1.06 | 0.13 | -0.07 | -0.64 | -0.21 | -0.80 | -0.31 | -0.76 | 0.00 |
| | zscore SD | 1.37 | 1.48 | 1.36 | 0.99 | 1.07 | 1.16 | 1.17 | 1.50 | 0.82 | 0.98 | 1.38 | 1.08 | 1.29 | 1.07 | 1.34 | 1.00 |
| | p-value (contrasted to controls) | 7.11E-01 | 8.81E-01 | 9.28E-01 | 9.97E-01 | 1.00E+00 | 9.60E-01 | 8.99E-01 | 5.42E-01 | 1.00E+00 | 1.00E+00 | 8.96E-01 | 1.00E+00 | 5.63E-01 | 2.43E-01 | 2.47E-03 | - |
| Emotional SDQ subscale (teacher report) | N | 7 | 11 | 5 | 11 | 5 | 19 | 11 | 6 | 6 | 26 | 10 | 10 | 11 | 95 | 43 | 52 |
| | orginal score | 3.29 | 3.27 | 4.00 | 2.45 | 1.40 | 3.00 | 2.36 | 2.17 | 2.83 | 2.23 | 4.20 | 3.50 | 2.91 | 2.63 | 3.33 | 1.83 |
| | orginal score SD | 2.98 | 2.10 | 3.00 | 2.30 | 1.14 | 2.47 | 1.12 | 1.72 | 2.23 | 1.75 | 2.20 | 2.55 | 3.18 | 2.11 | 2.49 | 2.15 |
| | zscore | -0.67 | -0.80 | -1.03 | -0.38 | 0.09 | -0.56 | -0.38 | -0.30 | -0.54 | -0.29 | -1.10 | -0.80 | -0.40 | -0.44 | -0.72 | 0.00 |
| | zscore SD | 1.13 | 0.94 | 1.20 | 0.94 | 0.74 | 1.12 | 0.55 | 0.94 | 1.07 | 0.89 | 0.72 | 0.96 | 1.32 | 0.93 | 1.04 | 1.00 |
| | p-value (contrasted to controls) | 8.92E-01 | 3.87E-01 | 5.25E-01 | 9.95E-01 | 1.00E+00 | 6.69E-01 | 9.97E-01 | 1.00E+00 | 9.88E-01 | 9.88E-01 | 7.30E-02 | 4.65E-01 | 9.88E-01 | 1.93E-02 | 7.01E-04 | - |
| Hyperactivity SDQ subscale (teacher report) | N | 7 | 11 | 5 | 11 | 5 | 19 | 11 | 6 | 6 | 26 | 10 | 10 | 11 | 95 | 43 | 52 |
| | orginal score | 4.29 | 2.55 | 5.40 | 3.73 | 4.00 | 4.26 | 3.91 | 5.67 | 3.17 | 3.15 | 4.60 | 3.20 | 5.73 | 3.66 | 4.60 | 1.27 |
| | orginal score SD | 2.14 | 2.81 | 2.41 | 2.57 | 2.74 | 3.05 | 2.84 | 1.97 | 1.83 | 1.89 | 2.55 | 2.44 | 2.15 | 2.43 | 2.66 | 1.74 |
| | zscore | -1.34 | -0.47 | -1.86 | -1.04 | -1.65 | -1.52 | -1.20 | -2.10 | -0.97 | -0.98 | -1.92 | -1.27 | -2.40 | -1.21 | -1.69 | 0.00 |
| | zscore SD | 1.67 | 1.90 | 1.05 | 1.65 | 1.13 | 1.66 | 1.36 | 1.47 | 0.78 | 1.12 | 1.32 | 1.27 | 1.46 | 1.35 | 1.63 | 1.00 |
| | p-value (contrasted to controls) | 2.10E-01 | 9.96E-01 | 6.25E-02 | 2.95E-01 | 2.80E-01 | 5.54E-04 | 1.04E-01 | 5.49E-03 | 8.35E-01 | 6.08E-02 | 8.90E-04 | 2.33E-01 | 6.34E-07 | 1.17E-07 | 5.93E-10 | - |
| Peer SDQ subscale (teacher report) | N | 7 | 11 | 5 | 11 | 5 | 19 | 11 | 6 | 6 | 26 | 10 | 10 | 11 | 95 | 43 | 52 |
| | orginal score | 3.57 | 2.82 | 1.80 | 3.18 | 3.60 | 3.26 | 2.82 | 3.50 | 3.00 | 3.08 | 3.20 | 1.50 | 3.09 | 2.99 | 2.95 | 1.29 |
| | orginal score SD | 2.57 | 2.82 | 1.64 | 2.40 | 2.97 | 2.31 | 2.71 | 1.22 | 2.37 | 2.24 | 1.81 | 1.08 | 2.26 | 2.29 | 2.12 | 1.84 |
| | zscore | -1.07 | -0.73 | -0.15 | -0.92 | -1.24 | -1.00 | -0.66 | -1.15 | -0.89 | -0.95 | -1.07 | -0.23 | -0.90 | -0.87 | -0.84 | 0.00 |
| | zscore SD | 1.40 | 1.48 | 0.86 | 1.23 | 1.52 | 1.13 | 1.39 | 0.58 | 1.36 | 1.19 | 0.94 | 0.75 | 1.34 | 1.19 | 1.16 | 1.00 |
| | p-value (contrasted to controls) | 4.00E-01 | 7.49E-01 | 1.00E+00 | 3.50E-01 | 5.30E-01 | 4.06E-02 | 8.10E-01 | 4.05E-01 | 8.20E-01 | 2.42E-02 | 2.30E-01 | 1.00E+00 | 3.91E-01 | 1.75E-05 | 6.07E-04 | - |
| Prosocial SDQ subscale (teacher report) | N | 7 | 11 | 5 | 11 | 5 | 19 | 11 | 6 | 6 | 26 | 10 | 10 | 11 | 95 | 43 | 52 |
| | orginal score | 5.43 | 6.27 | 4.80 | 4.91 | 4.80 | 4.74 | 5.91 | 4.00 | 5.83 | 5.35 | 5.30 | 7.20 | 3.73 | 5.44 | 4.91 | 7.65 |
| | orginal score SD | 3.46 | 3.41 | 2.17 | 2.84 | 4.09 | 2.66 | 3.24 | 2.10 | 2.79 | 3.03 | 2.26 | 2.66 | 3.04 | 2.98 | 2.84 | 2.09 |
| | zscore | -0.73 | -0.54 | -1.07 | -1.08 | -1.36 | -1.20 | -0.50 | -1.45 | -0.73 | -1.01 | -1.01 | -0.27 | -1.58 | -0.90 | -1.10 | 0.00 |
| | zscore SD | 1.53 | 1.54 | 0.92 | 1.49 | 2.29 | 1.25 | 1.62 | 1.12 | 1.23 | 1.35 | 0.99 | 1.11 | 1.44 | 1.40 | 1.29 | 1.00 |
| | p-value (contrasted to controls) | 9.36E-01 | 9.81E-01 | 7.70E-01 | 2.57E-01 | 5.71E-01 | 1.74E-02 | 9.81E-01 | 2.07E-01 | 9.78E-01 | 4.28E-02 | 4.84E-01 | 1.00E+00 | 7.84E-03 | 7.42E-05 | 4.48E-05 | - |

Supplementary Table 2



| Domain | ND-CNV Carriers | | | | | Controls | | | | | Group difference | | | | |
|---|---|---|---|---|---|---|---|---|---|---|---|---|---|---|---|
| | N | zscore | Lower CI | Upper CI | SD | N | zscore | Lower CI | Upper CI | SD | p | Cohen's d | Lower CI | Upper CI | Descriptor |
| FSIQ | 237 | -1.52 | -1.67 | -1.37 | 1.20 | 75 | 0.00 | -0.23 | 0.23 | 1.00 | <1.00E-15 | 1.32 | 1.03 | 1.60 | Large |
| PIQ | 238 | -1.06 | -1.20 | -0.93 | 1.07 | 75 | 0.00 | -0.23 | 0.23 | 1.00 | <1.00E-15 | 1.01 | 0.73 | 1.28 | Large |
| VIQ | 239 | -1.55 | -1.71 | -1.40 | 1.22 | 75 | 0.00 | -0.23 | 0.23 | 1.00 | <1.00E-15 | 1.33 | 1.04 | 1.61 | Large |
| Non-verbal reasoning | 238 | -0.98 | -1.13 | -0.83 | 1.15 | 75 | 0.00 | -0.23 | 0.23 | 1.00 | 2.50E-12 | 0.88 | 0.61 | 1.15 | Large |
| Perceptual organisation | 237 | -0.89 | -1.01 | -0.76 | 0.98 | 75 | 0.00 | -0.23 | 0.23 | 1.00 | 7.51E-13 | 0.90 | 0.63 | 1.18 | Large |
| Verbal knowledge | 237 | -1.33 | -1.47 | -1.19 | 1.08 | 75 | 0.00 | -0.23 | 0.23 | 1.00 | <1.00E-15 | 1.26 | 0.97 | 1.54 | Large |
| Verbal reasoning | 238 | -1.47 | -1.64 | -1.29 | 1.39 | 75 | 0.00 | -0.23 | 0.23 | 1.00 | <1.00E-15 | 1.13 | 0.85 | 1.40 | Large |
| Set-shifting | 198 | -0.79 | -0.95 | -0.62 | 1.15 | 68 | 0.00 | -0.24 | 0.24 | 1.00 | 1.66E-09 | 0.71 | 0.42 | 0.99 | Medium |
| Spatial working memory | 212 | -0.86 | -1.00 | -0.71 | 1.06 | 48 | 0.00 | -0.29 | 0.29 | 1.00 | 2.95E-07 | 0.82 | 0.49 | 1.14 | Large |
| Spatial planning | 170 | -0.63 | -0.78 | -0.49 | 0.94 | 46 | 0.00 | -0.30 | 0.30 | 1.00 | 5.99E-05 | 0.67 | 0.33 | 1.00 | Medium |
| Sustained attention | 155 | -1.31 | -1.53 | -1.08 | 1.39 | 44 | 0.00 | -0.30 | 0.30 | 1.00 | 5.73E-09 | 1.00 | 0.64 | 1.35 | Large |
| Processing speed | 194 | -0.41 | -0.55 | -0.27 | 0.99 | 46 | 0.00 | -0.30 | 0.30 | 1.00 | 3.90E-02 | 0.41 | 0.09 | 0.73 | Small |
| Total CAPA symptom count | 233 | -1.32 | -1.48 | -1.17 | 1.20 | 61 | 0.00 | -0.26 | 0.26 | 1.00 | <1.00E-15 | 1.14 | 0.84 | 1.44 | Large |
| Anxiety subscale | 233 | -0.82 | -1.00 | -0.64 | 1.41 | 61 | 0.00 | -0.26 | 0.26 | 1.00 | 7.85E-06 | 0.62 | 0.33 | 0.90 | Medium |
| ADHD subscale | 233 | -1.40 | -1.57 | -1.22 | 1.34 | 61 | 0.00 | -0.26 | 0.26 | 1.00 | 3.75E-14 | 1.09 | 0.79 | 1.39 | Large |
| Mood subscale | 233 | -0.67 | -0.82 | -0.52 | 1.16 | 61 | 0.00 | -0.26 | 0.26 | 1.00 | 2.43E-05 | 0.60 | 0.31 | 0.88 | Medium |
| OCD subscale | 233 | -0.92 | -1.21 | -0.64 | 2.23 | 61 | 0.00 | -0.26 | 0.26 | 1.00 | 3.88E-04 | 0.45 | 0.17 | 0.74 | Small |
| ODD subscale | 233 | -0.81 | -0.96 | -0.66 | 1.17 | 61 | 0.00 | -0.26 | 0.26 | 1.00 | 2.69E-06 | 0.72 | 0.43 | 1.00 | Medium |
| Sleep subscale | 233 | -0.66 | -0.84 | -0.49 | 1.32 | 61 | 0.00 | -0.26 | 0.26 | 1.00 | 1.14E-04 | 0.53 | 0.24 | 0.81 | Medium |
| Subclinical psychotic experiences | 244 | -0.49 | -0.74 | -0.23 | 2.03 | 66 | 0.00 | -0.25 | 0.25 | 1.00 | 5.68E-02 | 0.26 | -0.01 | 0.53 | Small |
| General functioning | 138 | -1.38 | -1.55 | -1.22 | 0.95 | 42 | 0.00 | -0.31 | 0.31 | 1.00 | <1.00E-15 | 1.45 | 1.06 | 1.83 | Large |
| Social functioning | 128 | -1.60 | -1.78 | -1.43 | 1.00 | 42 | 0.00 | -0.31 | 0.31 | 1.00 | <1.00E-15 | 1.62 | 1.21 | 2.02 | Large |
| ASD traits | 227 | -2.51 | -2.71 | -2.31 | 1.51 | 65 | 0.00 | -0.25 | 0.25 | 1.00 | <1.00E-15 | 1.78 | 1.46 | 2.10 | Large |
| Motor coordination | 226 | -1.94 | -2.10 | -1.77 | 1.24 | 65 | 0.00 | -0.25 | 0.25 | 1.00 | <1.00E-15 | 1.63 | 1.31 | 1.94 | Large |
| SDQ total (caregiver) | 229 | -1.87 | -2.02 | -1.72 | 1.15 | 67 | 0.00 | -0.24 | 0.24 | 1.00 | <1.00E-15 | 1.68 | 1.36 | 1.99 | Large |
| Conduct subscale (caregiver) | 229 | -0.85 | -1.01 | -0.69 | 1.23 | 67 | 0.00 | -0.24 | 0.24 | 1.00 | 5.22E-07 | 0.72 | 0.44 | 1.00 | Medium |
| Emotional subscale (caregiver) | 229 | -1.00 | -1.15 | -0.85 | 1.16 | 67 | 0.00 | -0.24 | 0.24 | 1.00 | 8.70E-11 | 0.89 | 0.61 | 1.17 | Large |
| Hyperactivity subscale (caregiver) | 229 | -2.63 | -2.83 | -2.42 | 1.57 | 67 | 0.00 | -0.24 | 0.24 | 1.00 | <1.00E-15 | 1.81 | 1.49 | 2.12 | Large |
| Peer subscale (caregiver) | 229 | -1.48 | -1.63 | -1.33 | 1.17 | 67 | 0.00 | -0.24 | 0.24 | 1.00 | <1.00E-15 | 1.31 | 1.01 | 1.60 | Large |
| Prosocial subscale (caregiver) | 229 | -0.93 | -1.07 | -0.79 | 1.08 | 67 | 0.00 | -0.24 | 0.24 | 1.00 | 1.38E-10 | 0.88 | 0.60 | 1.16 | Large |
| SDQ total (teacher) | 138 | -1.08 | -1.25 | -0.92 | 0.99 | 42 | 0.00 | -0.31 | 0.31 | 1.00 | 2.20E-10 | 1.10 | 0.73 | 1.47 | Large |
| Conduct subscale (teacher) | 138 | -0.34 | -0.53 | -0.16 | 1.11 | 42 | 0.00 | -0.31 | 0.31 | 1.00 | 7.15E-02 | 0.32 | -0.03 | 0.67 | Small |
| Emotional subscale (teacher) | 138 | -0.63 | -0.80 | -0.46 | 1.00 | 42 | 0.00 | -0.31 | 0.31 | 1.00 | 3.03E-04 | 0.63 | 0.27 | 0.98 | Medium |
| Hyperactivity subscale (teacher) | 138 | -1.55 | -1.82 | -1.29 | 1.59 | 42 | 0.00 | -0.31 | 0.31 | 1.00 | 8.83E-10 | 1.06 | 0.69 | 1.43 | Large |
| Peer subscale (teacher) | 138 | -0.85 | -1.06 | -0.65 | 1.20 | 42 | 0.00 | -0.31 | 0.31 | 1.00 | 1.87E-05 | 0.74 | 0.39 | 1.10 | Medium |
| Prosocial subscale (teacher) | 138 | -0.94 | -1.16 | -0.71 | 1.32 | 42 | 0.00 | -0.31 | 0.31 | 1.00 | 1.65E-05 | 0.75 | 0.39 | 1.10 | Medium |

| Domain | Genotype | | Gender | | | Age | | |
|---|---|---|---|---|---|---|---|---|
| | Eta % | p | Eta % | Beta | p | Eta % | Beta | p |
| FSIQ | 20.75 | 9.02E-05 | 1.25 | -0.10 | 1.08E-01 | 2.84 | -0.17 | 1.60E-02 |
| PIQ | 21.04 | 4.20E-05 | 1.93 | -0.12 | 4.21E-02 | 4.15 | -0.21 | 3.09E-03 |
| VIQ | 15.56 | 5.41E-03 | 0.22 | -0.04 | 5.14E-01 | 1.26 | -0.12 | 1.23E-01 |
| Non-verbal reasoning | 20.09 | 9.59E-05 | 0.84 | -0.07 | 1.83E-01 | 5.77 | -0.25 | 5.69E-04 |
| Perceptual organisation | 14.26 | 7.88E-03 | 4.05 | -0.19 | 5.08E-03 | 2.31 | -0.16 | 3.35E-02 |
| Verbal knowledge | 13.46 | 2.38E-02 | 0.43 | -0.06 | 3.79E-01 | 0.29 | -0.06 | 4.71E-01 |
| Verbal reasoning | 12.80 | 3.14E-02 | 0.11 | -0.02 | 6.56E-01 | 1.44 | -0.12 | 1.06E-01 |
| Set-shifting | 9.21 | 3.24E-01 | 1.36 | -0.13 | 1.56E-01 | 5.00 | 0.24 | 7.01E-03 |
| Spatial working memory | 10.62 | 1.72E-01 | 2.01 | -0.13 | 7.61E-02 | 1.99 | -0.15 | 7.76E-02 |
| Spatial planning | 12.93 | 2.06E-01 | 1.03 | 0.12 | 2.61E-01 | 1.70 | -0.14 | 1.49E-01 |
| Sustained attention | 14.62 | 6.55E-02 | 0.87 | 0.08 | 2.63E-01 | 23.18 | 0.50 | 1.00E-07 |
| Processing speed | 18.82 | 6.46E-03 | 2.03 | -0.16 | 7.66E-02 | 2.50 | 0.16 | 4.97E-02 |
| Total CAPA symptom count | 18.81 | 3.07E-04 | 2.55 | 0.15 | 2.23E-02 | 0.00 | 0.01 | 9.81E-01 |
| Anxiety subscale | 17.21 | 2.00E-03 | 1.14 | 0.12 | 1.41E-01 | 0.16 | -0.04 | 5.81E-01 |
| ADHD subscale | 14.33 | 7.95E-03 | 1.31 | 0.09 | 1.09E-01 | 1.86 | 0.15 | 5.66E-02 |
| Mood subscale | 10.16 | 9.89E-02 | 0.57 | 0.09 | 3.01E-01 | 2.20 | -0.14 | 4.36E-02 |
| OCD subscale | 6.37 | 5.23E-01 | 2.10 | 0.17 | 5.75E-02 | 1.55 | -0.13 | 1.02E-01 |
| ODD subscale | 14.46 | 1.06E-02 | 0.39 | 0.06 | 3.89E-01 | 0.31 | -0.05 | 4.47E-01 |
| Sleep subscale | 4.92 | 7.13E-01 | 3.79 | 0.18 | 9.93E-03 | 0.67 | 0.10 | 2.73E-01 |
| Subclinical psychotic experiences | 9.70 | 1.43E-01 | 0.05 | 0.04 | 7.54E-01 | 2.25 | -0.15 | 4.54E-02 |
| General functioning | 16.46 | 8.24E-02 | 0.49 | 0.07 | 4.41E-01 | 0.06 | 0.01 | 7.85E-01 |
| Social functioning | 20.67 | 2.93E-02 | 0.72 | 0.08 | 3.60E-01 | 0.17 | 0.03 | 6.57E-01 |
| ASD traits | 14.60 | 4.74E-03 | 0.88 | 0.11 | 1.81E-01 | 0.58 | -0.07 | 2.75E-01 |
| Motor coordination | 9.61 | 1.02E-01 | 2.59 | 0.16 | 2.50E-02 | 0.29 | 0.05 | 4.54E-01 |
| SDQ total | 17.92 | 3.95E-04 | 0.05 | 0.03 | 7.50E-01 | 0.45 | -0.06 | 3.28E-01 |
| Conduct subscale | 18.28 | 2.43E-04 | 0.25 | -0.05 | 4.65E-01 | 0.01 | 0.00 | 8.75E-01 |
| Emotional subscale | 9.87 | 9.84E-02 | 0.44 | -0.06 | 3.56E-01 | 1.03 | -0.10 | 1.60E-01 |
| Hyperactivity subscale | 12.54 | 1.20E-02 | 2.74 | 0.14 | 1.65E-02 | 5.04 | 0.23 | 1.24E-03 |
| Peer subscale | 9.11 | 1.16E-01 | 0.03 | 0.05 | 8.09E-01 | 6.27 | -0.26 | 4.83E-04 |
| Prosocial subscale | 14.16 | 6.84E-03 | 1.82 | 0.13 | 5.59E-02 | 0.20 | 0.05 | 5.24E-01 |
